# Uncovering functional deterioration in the rhizosphere microbiome associated with wheat dwarfing

**DOI:** 10.1101/2025.03.16.643510

**Authors:** Monique E. Smith, Vanessa N. Kavamura, David Hughes, Rodrigo Mendes, George Lund, Ian Clark, Tim H. Mauchline

## Abstract

**Background:** One of the biggest developments of wheat domestication was the development of semi-dwarf cultivars that respond well to fertilisers and produce higher yields without lodging. Consequently, this change has also impacted the wheat microbiome, often resulting in reduced selection of taxa and a loss of network complexity in the rhizospheres of semi-dwarf cultivars. Given the importance of rhizosphere microbiomes for plant health and performance, it is imperative that we understand if and how these changes have affected their function. Here, we use shotgun metagenomics to classify the functional potential of prokaryote communities from the rhizospheres of tall and semi-dwarf cultivars to compare the impact of wheat dwarfing on rhizosphere microbiome functions.

**Results:** We found distinct taxonomic and functional differences between tall and semi-dwarf wheat rhizosphere communities and identified that semi-dwarf wheat microbiomes were less distinct from bulk soil communities. Of the 113 functional genes that were differentially abundant between tall and semi-dwarf cultivars, 95 % were depleted in semi-dwarf cultivars and 65 % of differentially abundant reads best mapped to genes involved in staurosporine biosynthesis (antibiotic product), plant cell wall degradation (microbial mediation of plant root architecture, overwintering energy source for microbes) and sphingolipid metabolism (signal bioactive molecules).

**Conclusions:** Overall, our findings indicate that green revolution breeding has developed wheat cultivars with a reduced rhizosphere effect. The consequences of this are likely detrimental to the development of microbiome-assisted agriculture which will require a strong rhizosphere selective environment for the establishment of a beneficial plant root microbiome. We believe our results are of striking importance and highlight that implementation of microbiome facilitated agriculture as part of a sustainable crop production strategy will require an overhaul of wheat breeding programmes to consider plant-microbe interactions, especially in the root environment.

## Introduction

As a result of the Green Revolution, we benefit from a huge increase in cereal grain production. This was largely due to the incorporation of reduced-height (*Rht*) mutant allele dwarfing genes into modern wheat cultivars (i.e., wheat dwarfing). Nearly all wheat grown today are semi-dwarf cultivars with increased tillering, larger seed heads producing higher yields, but on shorter stems to prevent lodging, and respond well to high fertilizer inputs. However, current high-yielding semi-dwarf crop varieties rely on unsustainable levels of agrochemical inputs, including synthetic fertilisers, which are environmentally harmful (Van Grinsven et al., 2013).

While tall heritage cultivars or landraces are mainly grown on marginal fields on organic farms (Morgounov et al., 2016; Ortman et al., 2023) they are still an important genetic resource for breeding programs (Escudero-Martinez and Bulgarelli, 2023; Lammerts van Bueren and Myers, 2012), especially since they show tolerance to extreme weather events and other stresses (Newton et al., 2011). Wheat domestication gave little to no consideration to belowground processes; as such, we are only beginning to understand how wheat dwarfing has impacted the interactions between roots and soil organisms six decades after the first semi-dwarf wheat was introduced into agriculture (Kavamura et al., 2020; Rossmann et al., 2020; Tkacz et al., 2020).

The rhizosphere, i.e., the interface between roots and soil, harbours a dynamic community of microorganisms. These communities play an important role in how plants function, ranging from beneficial effects, e.g., aiding nutrient acquisition, growth promotion and plant defences (Mauchline and Malone, 2017; Mendes et al., 2013; Schlaeppi and Bulgarelli, 2015), to harmful effects, e.g., pathogens such as *Gaeumannomyces tritici* causative agent of take-all disease in most cereals (Palma-Guerrero et al., 2021). The assembly of rhizosphere communities are largely determined by the exudates and structure of plant roots (Sasse et al., 2018; Vives-Peris et al., 2020) which the microbiome itself can modulate (Korenblum et al., 2020; Pandey et al., 2021), therefore it is not surprising that the domestication of wheat, including wheat dwarfing, has been shown to influence protist, bacterial, nematode and fungal rhizosphere communities (Kavamura et al., 2020; Rossmann et al., 2020; Tkacz et al., 2020). The general trend in these populations is that wheat domestication has reduced selection processes in the rhizosphere. This is demonstrated by the root microbiome of ancestral wheat harbouring more unique taxa and more complex microbiomes than the modern cultivars. Furthermore, modern cultivars can be enriched in fungal pathogens (Kinnunen-Grubb et al., 2020). Similar trends have also been found in the domestication of other cereals such as barley (Bulgarelli et al., 2015), soybean (Shi et al., 2019), rice (Sun et al., 2021) and durum wheat (Spor et al., 2020). While these findings have been important for our understanding of microbial community dynamics there is now a requirement to understand microbiome assembly and function for these resources to be harnessed in sustainable agriculture programmes with less dependence on fertilizer inputs (Busby et al., 2017; Gruet et al., 2022). Phylogenetic marker gene profiles can hint at the function of these communities (Kavamura et al., 2020). However, to obtain a comprehensive understanding of how wheat dwarfing has impacted microbiome function, there is a requirement for holistic shotgun sequencing methods to be deployed.

Whole shotgun metagenomic sequencing provides a representation of the genomes present in each sample, allowing the inference of functional potential of a microbial community. This technique has been used to study the wheat rhizosphere microbiome and identify microbes that consume plant-derived carbon (Fan et al., 2022), to ascertain differences in microbial zinc-mobilisation genes between high and low zinc wheat cultivars (Wang et al., 2021a), and the comparison of antibiotic resistance genes in the rhizospheres of common crops (Yu et al., 2023). To date, shotgun metagenomics has not corroborated the wheat domestication-driven changes in rhizosphere communities found using phylogenetic marker gene studies. A previous attempt was made in Canadian wheat cultivars and found no effect of domestication, but this study sampled at wheat senescence (Quiza et al., 2021), a time when the structure of the root microbiome of annual plants has been found to degrade and become dominated by saprophytes (Mauchline et al., 2015). However, a similar study conducted in durum (tetraploid) wheat demonstrated that domestication led to a decline in gene diversity and a shift in microbial functional traits, particularly related to nutrient cycling (Yue et al., 2023). As yet, the impact of wheat dwarfing on the vegetative stages of hexaploid bread wheat plants has not yet been explored using shotgun metagenomics.

Given the strong evidence from previous studies that wheat dwarfing impacts the rhizosphere community structure at a taxonomic level, we predict that the same impact will be observed for a range of functional genes in these communities. We tested this hypothesis by growing two tall and two semi-dwarf wheat cultivars alongside unplanted bulk soil control pots under glasshouse conditions. At flowering stage, we sampled the rhizosphere soil and bulk soil samples and generated, analysed, and compared the shotgun metagenomic profiles, in terms of taxonomy and function, of the prokaryote communities from these sample types.

## Methods

### Wheat cultivars and growth conditions

Wheat cultivars (Table 1) were selected based on previous results by Kavamura *et al*., 2020, which indicated differences in the root microbiome structure and predicted function between short and tall cultivars. Soil was collected from Stackyard bare-fallow soil mine (52.002997°N, 0.613058°W) in January 2019. Soil details are previously described (Reid et al., 2021). Soil was sieved (2 mm mesh), mixed thoroughly, and stored at 4°C in polythene bags prior to use. Seeds were surface sterilized as previously described (Robinson et al., 2016) and were transferred to pre-soaked (sterile distilled water) filter paper in Petri dishes and germinated for three days in the dark at room temperature. Seedlings were transplanted to individual wells on a seed tray (1x seedling per well) in Stackyard bare fallow soil and grown in a glasshouse at Rothamsted Research for two weeks (20 °C, 16 h/day light regime, watered daily) before vernalization for twelve weeks (4 °C; 8 h light and 16 h dark). After this time, 3 seedlings were transplanted to 6-inch diameter pots (approximately 1 kg soil per pot), with NPK granules [15% N, 9% P_2_O_5_, 11% K_2_O, 2% MgO with micro-nutrients (B, Cu, Fe, Mn, Mo, and Zn); Osmocote, United Kingdom] (∼5 g per pot) added to the soil surface of each pot. Five replicate pots were prepared for each wheat variety and three unplanted bulk soil control pots were also set up using the same soil and fertilisation. Plants were grown in a glasshouse (20°C, 16 h/day light regime) and watered daily or as required with tap water.

**Table 1.** Characteristics of the cultivars used in this study.

| Cultivar | Year | Pedigree | Height |
| --- | --- | --- | --- |
| Chidham White Chaff | 1790 | Not recorded | Tall |
| Red Lammas | 1850 | Not recorded | Tall |
| Malacca | 1997 | Riband*(Rendezvous)*Apostle | Semi-dwarf |
| Gallant | 2009 | (Malacca*Charger)*Xi-19 | Semi-dwarf |

\*Table was modified from Kavamura *et al.* (2020).

Pots were harvested at the start of flowering (Zadoks growth stage 61; approximately 10 weeks growth post vernalisation; Zadoks et al., 1974), resulting in twenty rhizosphere samples and three bulk soil samples. Loose soil from the root system of a given pot was carefully removed. A 10 g subsample of root system was transferred to a 50 ml Falcon tube and 30 ml sterile water added. Next, samples were shaken vigorously for 10 min using an orbital shaker to release rhizosphere soil. After this time 4 ml soil suspension was centrifuged (2 min, RT, 15,000 rpm), supernatant discarded, and remaining soil was flash frozen in liquid nitrogen and stored at −80 °C.

### DNA extraction and sequencing

Genomic DNA was extracted from the bulk soil and rhizosphere soil sample (∼0.25 g) using the Dneasy PowerSoil Pro kit (Qiagen, Venlo, Netherlands) and stored at −80 °C. Extractions were performed according to the manufacturer’s instructions and with the use of the MP Biomedicals FastPrep-24 machine twice for the bead-beating step at 30 s at 5.5 m s^-1^. DNA purity and concentrations were measured with a NanoDrop 1000 spectrophotometer (Thermo Fisher Scientific, Wilmington, DE, United States), as well as a Qubit 2.0 Fluorimeter using the dsDNA HS assay kit (Thermo Fisher Scientific). 0.6 µg of DNA for each sample was sent to Novogene (UK) Company Limited for library preparation and sequencing using Illumina NovaSeq 6000 platform (HWI-ST1276) using a 150bp paired-end sequencing strategy. An average of 2.65^8^ raw paired reads, ranging between 1.66^8^ – 3.46^8^) per sample was obtained. Reads were quality checked using the FASTX-Toolkit (http://hannonlab.cshl.edu/fastx_toolkit/index.html; v0.0.14) with a quality threshold of 25, and Trimmomatic (http://www.usadellab.org/cms/?page=trimmomatic; v0.39; Bolger et al., 2014) with a minimum length of 80 bp. Quality checked reads were assigned to taxa using DIAMOND (https://github.com/bbuchfink/diamond; v2.0.13; Buchfink et al., 2015) and AnnoTree (http://annotree.uwaterloo.ca/annotree/app/downloads.html; v1.2; Gautam et al., 2022; Mendler et al., 2019) for prokaryote identification. KEGG Orthology (KO) molecular functional identifiers were assigned by MEGAN6 Ultra (https://software-ab.cs.uni-tuebingen.de/download/megan6/welcome.html; v6.24.23; Bağcı et al., 2019; Huson et al., 2016) and used to extract individual gene sequences for each functional KEGG identifier. Thus, processing resulted in two abundance tables for further analysis, one where reads were aligned to prokaryote taxa and another to functional genes.

### Statistical analysis

Taxa and functional genes were removed if their total count was < 10 reads. Only taxa assigned to Phyla or lower were kept for further analysis and reads unclassified to functional genes were removed. This resulted in 5,908,371,442 reads for the taxonomy table and 4,392,129,033 for the functional table and in both cases samples with the most reads roughly double those with the lowest number of reads.

All statistical analysis and visualisation of results was done in R version 4.3.3 (R core Team, 2024). Alpha and beta diversities were calculated from rarefied data (Weiss et al., 2017) while differential abundance analysis was done using DESeq2 variance stabilisation technique (McMurdie and Holmes, 2014) to normalise taxonomy and function abundance tables. The rarefied tables were calculated by normalizing sequence number to minimum sample size (159,120,960 and 118,292,681 for taxonomy and function tables respectively) by random subsampling without replacement using *rarefy_even_depth* function in the phyloseq package version 1.46 (McMurdie and Holmes, 2013). Rarefaction curve analysis was used to test that the subsampling of sequences still yielded sufficient resolution of prokaryote communities and their functional genes (Fig. S1).

To test whether the sample type, including tall wheat rhizosphere, semi-dwarf wheat rhizosphere and bulk soil, impacted alpha diversity, we obtained observed and Chao1 richness and Shannon diversity from the rarefied taxonomy and functional tables using *estimate_richness* function in the phyloseq package. Normality and homogeneity of variances of alpha diversity measures were tested before performing type 3 one-way ANOVA to account for unbalanced design due to fewer replicates in the bulk soil samples, and Tukey’s honest significant differences (HSD) with sample type as a main factor.

#### Beta diversity

Sample differences of the rarefied taxonomy and functional tables were visualised with principal coordinate analysis (PcoA) plots using Bray Curtis dissimilarity. To statistically test for differences between sample types we used Permutational Multivariate Analysis of Variance (PERMANOVA) with Bray Curtis distance matrices using *adonis2* in the vegan package version 2.6-4 (Oksanen et al., 2024) with 9,999 permutations. Pairwise comparisons of each group were evaluated with *pairwise.adonis* using a false discovery rate correction for multiple tests. We evaluated differences among sample variability using homogeneity of multivariate dispersions tests (*betadisper*), followed by ANOVAs to compare the mean distance-to-centroid. Pairwise dispersion comparisons were carried out using Tukey’s HSD.

#### Differential abundance analysis

Differential abundances of individual 21,805 taxa and 10,189 functional genes were calculated using DESeq2 package version 1.42.1 (Love et al., 2014) which is particularly powerful for small datasets (Li et al., 2022). Maximum-likelihood estimates for the log2-fold change between conditions associated with each gene or taxa were calculated using a negative binomial generalized linear model. Contrasts between each level of sample type were made and tests of significance were conducted using Wald’s test, employing α[=[0.05 and a Benjamin–Hochberg false discovery rate (q) of 0.05 to control type I error rate in the face of multiple comparisons. Bayesian adaptive shrinkage was then applied to reduce the log2-fold change towards zero for taxa or genes with low mean counts or a high dispersion in their count distribution (Stephens, 2017). We then considered taxa or genes to be enriched in a particular sample type only if the resulting shrunken log2-fold changes were > 1 or < −1, i.e. double in abundance (Sarno et al., 2023; Semaan et al., 2023).

Differential abundance analysis was done on all genes but for visualisation we only showed those that were considered enriched in a particular sample type and we compared the results of the analysis from these against ten housekeeping genes used as a baseline. These housekeeping genes were selected to cover different parts of the genome and that encode proteins involved in different metabolic activity, except for the ribosomal genes, similar to the gene selection in multilocus sequence typing (MLST; Takle et al., 2007; Maiden et al., 2013). These included signal recognition particle protein (*ffh*), glutamine synthetase (*glnA*), DNA gyrase (*gyrB*), transcription termination factor Rho (*rho*), 50S ribosomal protein L9 (*rplI*), 50S ribosomal subunit protein L17 (*rplQ*), RNA polymerase (*rpoZ),* DNA topoisomerase I (*topA*), nucleoside-specific channel-forming protein (*tsx*), ATP synthase F1, β-subunit (*atpD*).

## Results

### Alpha diversity

Overall, alpha diversity differences were statistically significant between sample types, i.e., bulk soil and rhizosphere soil from semi-dwarf and tall cultivars, for both taxonomy and function of the prokaryote communities (Fig. 1A, C). Tall cultivars harboured fewer taxa (observed and chao1) in their rhizospheres than bulk soils and semi-dwarf rhizospheres; however, when rarefied read abundances were considered, their communities were the most diverse (Shannon’s species diversity). Functional genes followed a similar pattern, though the differences were less profound. Shannon’s diversity was marginally greater in tall rhizospheres and more genes ascribed to semi-dwarf than tall rhizospheres. Wheat rhizospheres regardless of cultivar were found to have a higher functional diversity than bulk soil samples.

**Fig. 1.**
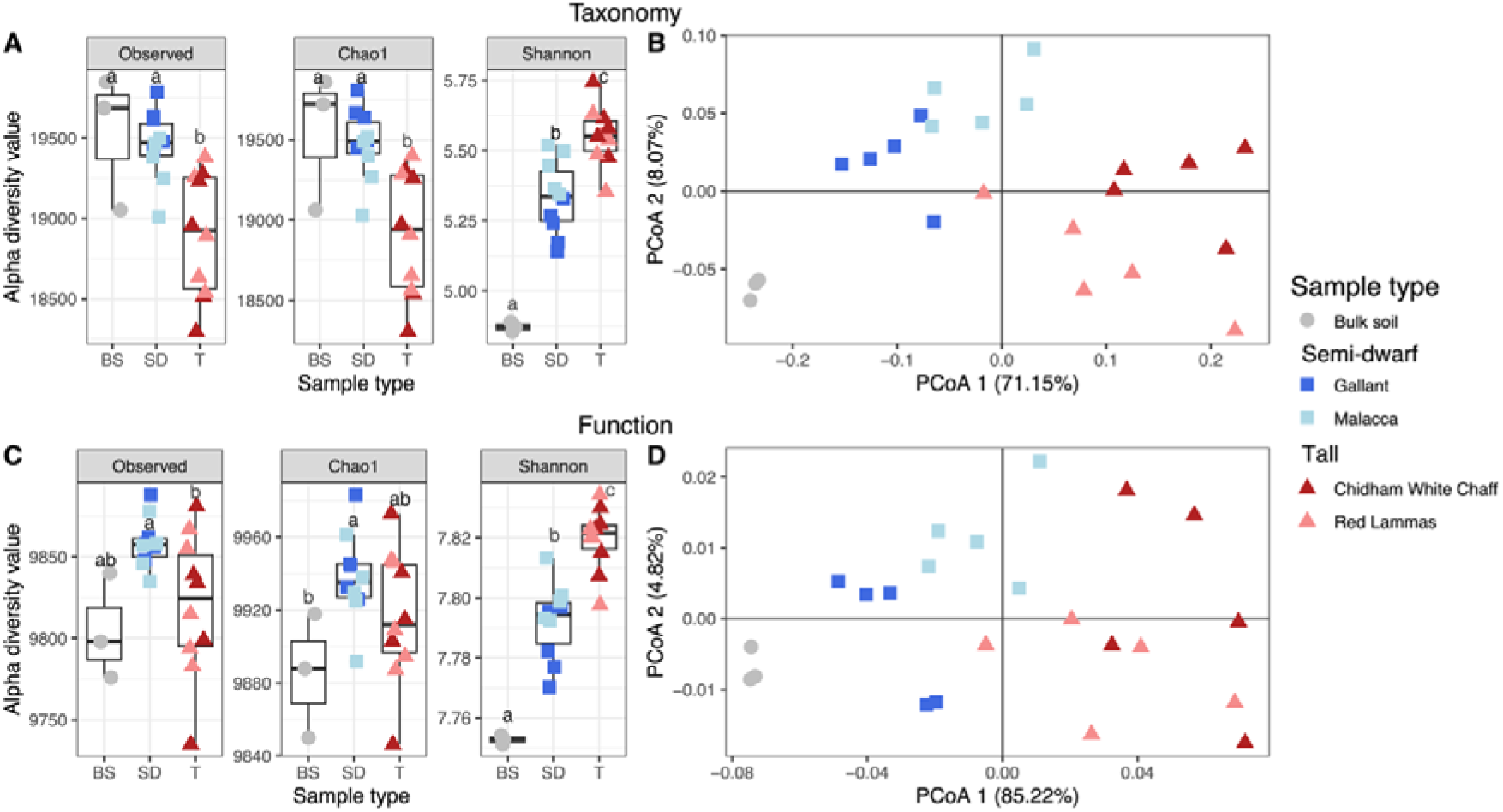
Diversity measures of the taxonomy and function of prokaryote communities. Alpha diversity is represented in plots A and C, and Principal Coordinate Analysis (PCoA) plots based on Bray-Curtis dissimilarity demonstrate beta diversity in plots B and D, for taxonomy and function respectively. Significant differences are shown between sample types, bulk soil samples (circles) and the rhizospheres of semi-dwarf (squares) and tall (triangles) wheat. Colours differentiate between the wheat different cultivars.

### Beta diversity

We found very similar PERMANOVA results for both taxonomy and function-assigned genes, whereby a high proportion of variation between samples was explained by sample type (bulk soil, semi-dwarf and tall wheat rhizospheres; Table 2). Pairwise comparisons between sample types were all significantly different with the highest R^2^ between tall rhizosphere and bulk soils. This difference is partly explained by significantly different dispersions between groups for taxonomy and function (ANOVA, F = 11.9, *p* < 0.001; F = 6.8, *p* = 0.006, respectively). *Posthoc* comparisons revealed that all groups differed in dispersion for taxonomy, but for function, there was less dispersion in the bulk soil samples compared to semi-dwarf (*padj* = 0.056) and tall (*padj* = 0.004) rhizospheres. Nevertheless, the PCoA clearly supports the PERMANOVA results that sample type is an important factor in determining the prokaryote taxonomy and function since they were clustered separately in multivariate space (Fig. 1B, D). Most of the variation was explained by PCoA 1 for both taxonomy and functional datasets which demonstrated that, from the cultivars included here, semi-dwarf wheat cultivars harbour prokaryote community profiles that are more similar to bulk soil than tall cultivars do. The shift in the prokaryote community was represented by an increasing relative abundance of Pseudomonadota (Proteobacteria) and Bacteroidota from bulk soil to rhizosphere soil of semi-dwarf and tall wheats and a higher proportion of Acidobacteroidota in bulk soil samples (Fig. S2).

**Table 2.**
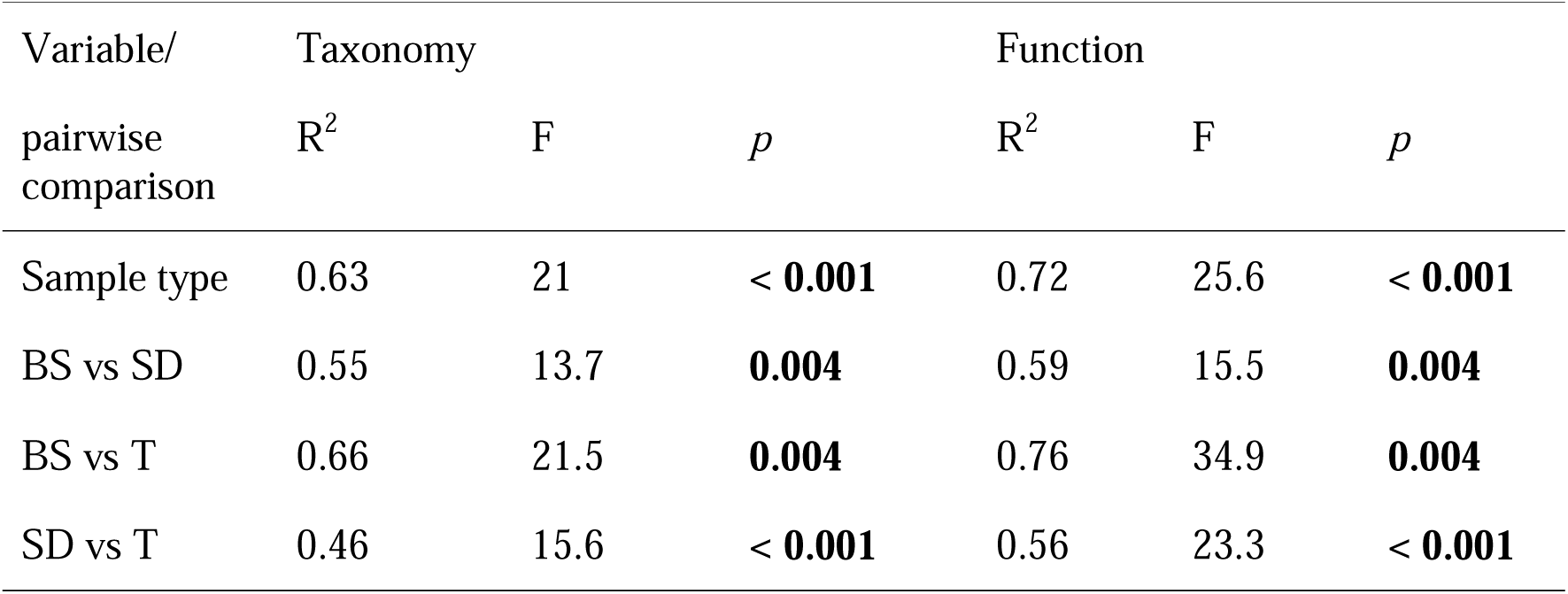
PERMANOVA and Tukey’s pairwise results for both taxonomy and function of Prokaryote communities. All combinations of sample type, bulk soil (BS) and rhizosphere soil of semi-dwarf (SD) and tall (T) cultivars, were present. Significant differences in bold and *p* values of pairwise tests have been adjusted for multiple comparisons using a false-discovery rate correction.

### Differential abundance

Out of 21,805 taxa and 10,189 functional genes, 5,072 (23 %) and 1,719 (17 %) respectively were differentially abundant for at least one contrast between the sample types (Fig. 2; Ward *p* < 0.05, FDR < 0.05, shrunken log2 fold > 1 or < −1). These differentially abundant taxa and functional genes make up 46 % and 5 % of total raw reads, respectively. The primary aim of this work is to compare abundances of genes between wheat genotypes. However, the biggest contrast was between tall rhizosphere and bulk soil prokaryote communities whereby 4,975 taxa and 1,712 functional genes were differentially abundant. Semi-dwarf rhizosphere communities were more similar to bulk soil communities than the tall rhizosphere communities with 1,114 taxa and 194 functional genes differentially abundant, 78 % and 89 % less than the tall vs bulk soil contrast, respectively. Despite this difference, the taxa that were enriched in either the tall or semi-dwarf wheat rhizosphere samples, compared with the bulk soil, belonged mostly to the same three phyla, Pseudomonadota, Bacteroidota and Actinomycetota (Actinobacteria), which when combined, accounted for 95 % and 92 % of the enriched taxa respectively (Fig. S3). Bulk soil samples were enriched in taxa from a wide range of phyla when compared to both rhizosphere types, dominated by Bacillota (Firmicutes), but the highest proportion of taxa belonging to various rare phyla (phyla that contain less than 5 % of all enriched taxa). Interestingly, when assessing the abundances of the differentially abundant functional genes between tall rhizospheres and bulk soil, 19 of the 20 most abundant genes were also more abundant in tall than semi-dwarf rhizospheres (ANOVA, *p* < 0.05; Table S1). The one exception being K07305 (peptide-methionine (R)-S-oxide reductase) which was equally abundant in the rhizosphere of tall and semi-dwarf but has low representation in bulk soil samples. It was found that 13 genes out of these 20 most abundant genes were also more abundant in semi-dwarf rhizospheres compared to bulk soil samples.

**Fig. 2.**
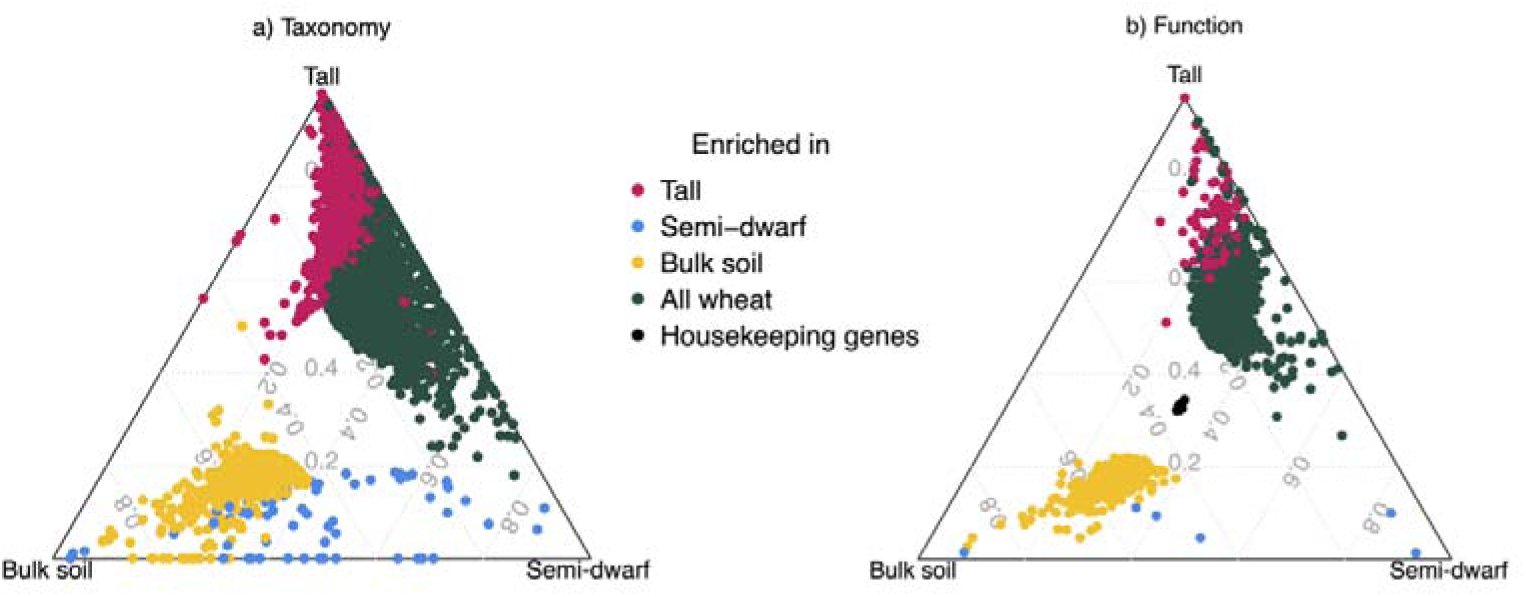
Ternary plots of differentially abundant taxonomy (a) and function (b) of prokaryote communities. Each point represents a taxa (a) or functional gene (b) and their position represents the contribution of the indicated sample type, bulk soil, semi-dwarf and tall rhizospheres, to the total normalised abundance. Taxa or functional genes are significantly different in abundance for different contrasts (Ward *p* < 0.05, FDR < 0.05, shrunken log fold changes > 1) and their colours indicate the group they are enriched in (i.e., higher abundance). Taxa or functional genes enriched in ‘all wheat’ samples, i.e., rhizosphere samples regardless of cultivar type, were determined by contrasts with bulk soil samples and vice versa for enriched in bulk soil. Enriched in tall rhizospheres was determined compared with semi-dwarf rhizospheres, regardless of bulk soil and vice versa for semi-dwarf. Therefore, some points could be characterised by two colours but the comparison between tall and semi-dwarf rhizospheres is the one presented as this is the most relevant for our hypothesis. Prokaryote housekeeping genes were included in b as a reference for differentially abundant functional genes (see methods for a list of housekeeping genes).

There were 1,414 taxa and 113 functional genes that were differentially abundant between the semi-dwarf and tall rhizosphere communities, most of which (95 %) were enriched in the tall rhizosphere communities (Fig. 3). The taxa enriched in the tall rhizosphere communities followed the same pattern as when compared to bulk soil whereby 96 % of reads were from Pseudomonadota, Bacteroidota and Actinomycetota (Fig. S3). The semi-dwarf rhizosphere communities were mostly enriched in Bacillota, Pseudomonadota, Spirochaetota and the highest proportion of enriched taxa belonged to rare phyla (phyla that contain less than 5 % of all enriched taxa). The differentially abundant functional genes belong to 15 different functional groups whereby tall rhizospheres had enriched genes belonging to each group (6,574,723 raw reads) and the six functional genes enriched in the semi-dwarf rhizospheres belonged to three categories (3,257 raw reads; Fig. 3b). Of the 107 genes enriched in the rhizospheres of tall wheats, 40% of reads mapped to genes associated with secondary metabolism, 21% with plant cell wall degradation, 10% with two-component sensor regulation, 8% with membrane transport, 12% with primary metabolism (other than cell wall degradation), 2% with secretion systems, 2% to transcriptional regulation and the remaining 5% to quorum sensing, cell cycle, biofilm formation, antibiotic resistance, motility, defence, and secondary messaging functions (Table 3).

**Fig. 3.**
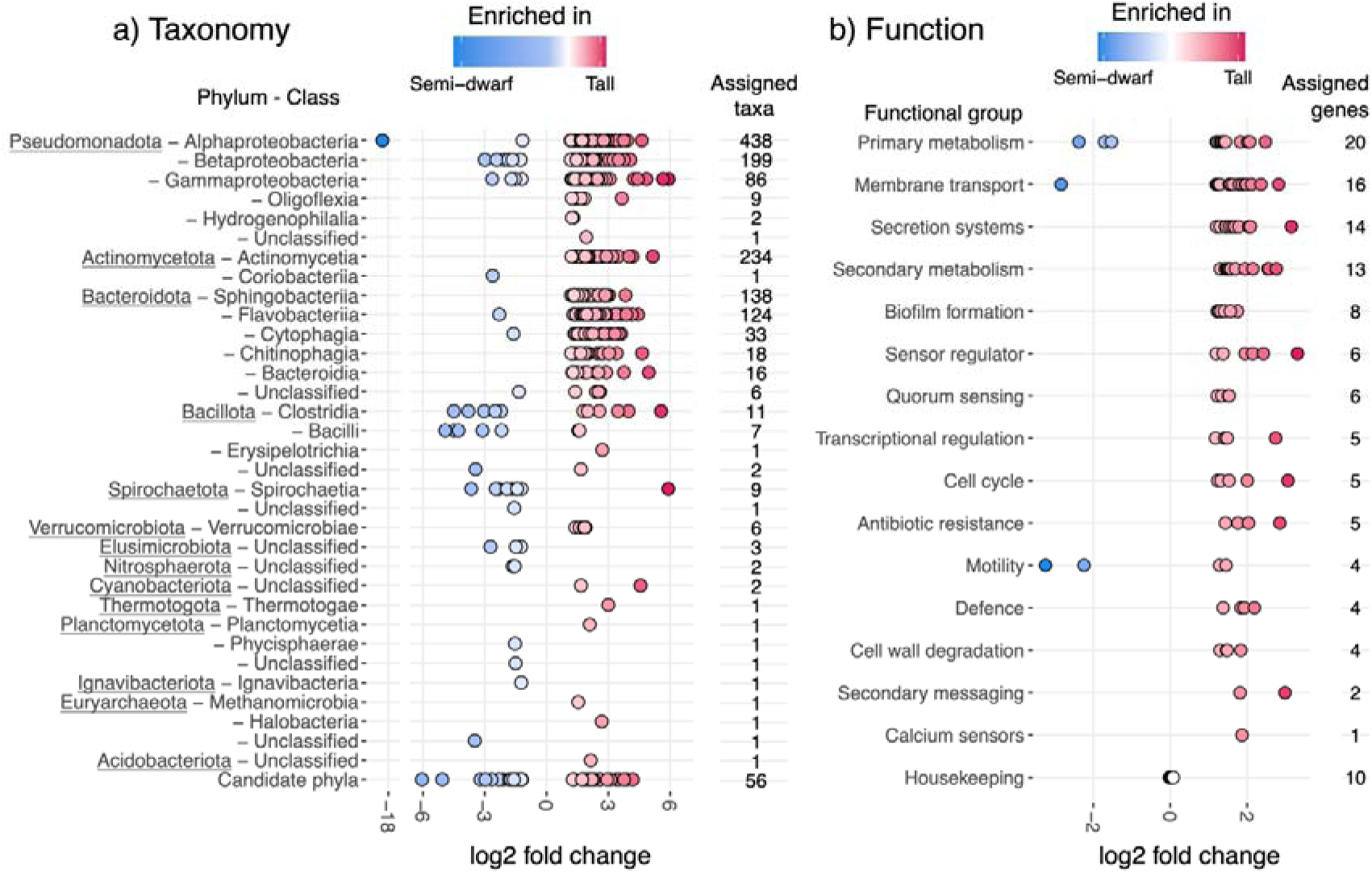
Differentially abundant taxonomy (a) and function (b) of prokaryote communities between semi-dwarf and tall wheat rhizospheres. Taxa and functional genes have been assigned to Class or functional group (based on KEGG orthology) respectively and the number of differentially abundant taxa or functional genes in each group are listed to the left of each plot. Only significant (Ward *p* < 0.05, FDR < 0.05,) and large (at least double; shrunken log2 fold > 1 or < −1) differences are shown except for the 10 housekeeping genes included for reference against the differentially abundant functional genes.

**Table 3.** Summaries of the taxa assigned to the 113 functional genes identified as differentially abundant between tall and semi-dwarf wheat rhizospheres. The functional genes are identified by the Kyoto Encyclopaedia of Genes and Genomes identifier (KEGG ID) and put into broad functional groups. The total read count refers to the total number of reads for each functional gene found in bulk soil and rhizosphere samples normalised by DESeq2 size factors for each sample (ref). The proportion of reads belonging to either semi-dwarf or tall rhizosphere samples are indicated, and genes enriched in the tall and semi-dwarf rhizospheres are separated by the horizontal line. Reads were reassigned to the taxa containing that gene and richness was calculated as the observed richness, i.e., the total number of taxa in either tall or semi-dwarf samples containing the corresponding functional gene, and Shannon’s diversity is the richness weighted by normalised read count to consider the evenness of the community. Numbers in bold represent significant differences between the means of either richness or diversity between semi-dwarf and tall rhizosphere samples (n = 20) as determined by Welch Two Sample t-tests (*p* < 0.05) using a false-discovery rate correction to adjust for multiple comparisons. *NOTE:* Table 3 *exceeds one A4 page so has been included as an additional file for reviewing as per journal requirements*.

| KEGG ID | Function | Gene name | Total read count<br>(normalised) | Proportion of read count | | Observed richness<br>(mean±sd) | | Shannons diversity<br>(mean±sd) | |
| --- | --- | --- | --- | --- | --- | --- | --- | --- | --- |
|  |  |  |  | Semi-dwarf | Tall | Semi-dwarf | Tall | Semi-dwarf | Tall |
| K14266 | Secondary metabolism | tryptophan 7-halogenase [EC:1.14.19.9] | 2417360 | 27 % | 71 % | **174 ± 19** | **243 ± 19** | **3.5 ± 0.1** | **3.7 ± 0.1** |
| K16397 | Secondary metabolism | epothilone polyketide synthase D | 29204 | 23 % | 72 % | 11 ± 2 | 11 ± 2 | **1.8 ± 0.1** | **1.5 ± 0.3** |
| K16403 | Secondary metabolism | O-methyltransferase | 4801 | 22 % | 74 % | 5 ± 1 | 6 ± 1 | **1.3 ± 0.2** | **1.0 ± 0.1** |
| K14627 | Secondary metabolism | dehydratase [EC:4.2.1.-] | 3362 | 22 % | 75 % | **5 ± 2** | **10 ± 2** | **1.3 ± 0.3** | **1.7 ± 0.2** |
| K14368 | Secondary metabolism | 3-alpha-mycarosylerythronolide B desosaminyl transferase [EC:2.4.1.278] | 2491 | 21 % | 74 % | **4 ± 1** | **7 ± 1** | 1.2 ± 0.3 | 1.4 ± 0.3 |
| K11009 | Secondary metabolism | murine toxin | 2377 | 21 % | 76 % | 3 ± 1 | 4 ± 1 | **0.9 ± 0.2** | **0.6 ± 0.4** |
| K19885 | Secondary metabolism | dichlorochromopyrrolate synthase / catalase [EC:1.21.98.2 1.11.1.6] | 1871 | 16 % | 80 % | **3 ± 1** | **5 ± 1** | 0.6 ± 0.4 | 0.9 ± 0.2 |
| K21212 | Secondary metabolism | NDP-hexose 2,3-dehydratase | 1851 | 20 % | 76 % | **3 ± 1** | **5 ± 1** | **0.8 ± 0.3** | **1.2 ± 0.3** |
| K16448 | Secondary metabolism | methylation protein MtfA | 1573 | 16 % | 81 % | **2 ± 1** | **5 ± 1** | **0.6 ± 0.4** | **1.1 ± 0.2** |
| K15968 | Secondary metabolism | tetracenomycin F2 cyclase [EC:4.2.1.154] | 1019 | 19 % | 76 % | **2 ± 1** | **4 ± 1** | **0.4 ± 0.4** | **1.1 ± 0.2** |
| K20086 | Secondary metabolism | tryptophan oxidase VioA [EC:1.4.3.-] | 203 | 8 % | 91 % | **0 ± 0** | **2 ± 1** | **0 ± 0** | **0.2 ± 0.2** |
| K20090 | Secondary metabolism | violacein synthase [EC:1.14.13.224] | 191 | 8 % | 92 % | **0 ± 0** | **1 ± 1** | 0 ± 0 | 0.1 ± 0.3 |
| K20088 | Secondary metabolism | violacein biosynthesis protein VioE | 125 | 10 % | 84 % | **0 ± 0** | **1 ± 1** | **0 ± 0** | **0.3 ± 0.3** |
| K18578 | Cell wall degradation | xyloglucan-specific exo-beta-1,4-glucanase [EC:3.2.1.155] | 609196 | 25 % | 73 % | **120 ± 12** | **146 ± 10** | **3.7 ± 0.1** | **3.6 ± 0.1** |
| K18651 | Cell wall degradation | oligoxyloglucan reducing-end-specific cellobiohydrolase [EC:3.2.1.150] | 537163 | 25 % | 73 % | **112 ± 11** | **131 ± 9** | 3.5 ± 0.0 | 3.5 ± 0.1 |
| K18786 | Cell wall degradation | cellobionic acid phosphorylase [EC:2.4.1.321] | 75143 | 20 % | 78 % | **18 ± 4** | **24 ± 3** | **2.3 ± 0.1** | **2.0 ± 0.1** |
| K18576 | Cell wall degradation | xyloglucan-specific endo-beta-1,4-glucanase [EC:3.2.1.151] | 69699 | 26 % | 70 % | **28 ± 4** | **40 ± 5** | **2.5 ± 0.1** | **2.7 ± 0.1** |
| K01202 | Primary metabolism | galactosylceramidase [EC:3.2.1.46] | 322927 | 27 % | 68 % | **66 ± 10** | **83 ± 5** | 3.1 ± 0.1 | 3.1 ± 0.1 |
| K20455 | Primary metabolism | 2-methylcitrate dehydratase (2-methyl-trans-aconitate forming) [EC:4.2.1.117] | 150825 | 27 % | 69 % | **24 ± 4** | **32 ± 3** | **2.1 ± 0.1** | **2.2 ± 0.1** |
| K02847 | Primary metabolism | O-antigen ligase [EC:2.4.1.-] | 101796 | 27 % | 70 % | **54 ± 7** | **79 ± 12** | 3.2 ± 0.1 | 3.2 ± 0.2 |
| K08961 | Primary metabolism | chondroitin-sulfate-ABC endolyase/exolyase [EC:4.2.2.20 4.2.2.21] | 45190 | 20 % | 77 % | 18 ± 3 | 20 ± 2 | 1.8 ± 0.1 | 1.9 ± 0.1 |
| K08325 | Primary metabolism | NADP-dependent alcohol dehydrogenase [EC:1.1.-.-] | 36347 | 26 % | 71 % | **21 ± 4** | **30 ± 3** | **2.5 ± 0.2** | **2.6 ± 0.1** |
| K00211 | Primary metabolism | prephenate dehydrogenase (NADP+) [EC:1.3.1.13] | 33022 | 25 % | 73 % | **7 ± 2** | **9 ± 2** | 0.6 ± 0.2 | 0.7 ± 0.2 |
| K03181 | Primary metabolism | chorismate lyase [EC:4.1.3.40] | 26153 | 27 % | 69 % | **22 ± 3** | **26 ± 5** | **2.6 ± 0.1** | **2.4 ± 0.2** |
| K01085 | Primary metabolism | glucose-1-phosphatase [EC:3.1.3.10] | 13928 | 24 % | 73 % | **13 ± 3** | **17 ± 2** | **2.1 ± 0.2** | **1.7 ± 0.4** |
| K01355 | Primary metabolism | omptin [EC:3.4.23.49] | 10723 | 25 % | 75 % | **7 ± 2** | **13 ± 2** | **1.0 ± 0.4** | **1.8 ± 0.3** |
| K01819 | Primary metabolism | galactose-6-phosphate isomerase [EC:5.3.1.26] | 8410 | 24 % | 72 % | **10 ± 1** | **13 ± 2** | 2.1 ± 0.1 | 2.0 ± 0.1 |
| K16215 | Primary metabolism | 2-ketoarginine methyltransferase [EC:2.1.1.243] | 4920 | 22 % | 73 % | 6 ± 1 | 7 ± 2 | 1.3 ± 0.3 | 1.0 ± 0.2 |
| K00998 | Primary metabolism | CDP-diacylglycerol---serine O-phosphatidyltransferase [EC:2.7.8.8] | 4027 | 24 % | 72 % | **4 ± 1** | **5 ± 1** | 1.1 ± 0.2 | 1.1 ± 0.2 |
| K21280 | Primary metabolism | 3-hydroxy-4-methylanthranilyl-[aryl-carrier protein] 5-monooxygenase [EC:1.14.13.223] | 3575 | 15 % | 84 % | 2 ± 1 | 2 ± 1 | **0.4 ± 0.3** | **0 ± 0** |
| K12455 | Primary metabolism | CDP-abequose synthase [EC:1.1.1.341] | 3208 | 17 % | 80 % | 5 ± 1 | 6 ± 1 | **1.4 ± 0.2** | **0.7 ± 0.2** |
| K21239 | Primary metabolism | virion DNA-directed RNA polymerase [EC:2.7.7.6] | 3016 | 18 % | 81 % | 2 ± 1 | 2 ± 0 | 0.2 ± 0.2 | 0.1 ± 0.1 |
| K19033 | Primary metabolism | 30S ribosomal protein S31 | 2812 | 27 % | 68 % | **6 ± 2** | **9 ± 1** | **1.4 ± 0.2** | **1.9 ± 0.1** |
| K03397 | Primary metabolism | indoleacetate---lysine synthetase<br>[EC:6.3.2.20] | 413 | 10 % | 88 % | **0 ± 0** | **1 ± 1** | 0 ± 0 | 0.1 ± 0.2 |
| K05874 | Sensor regulator | methyl-accepting chemotaxis protein I,<br>serine sensor receptor | 483244 | 25 % | 72 % | **108 ± 18** | **178 ± 22** | **3.4 ± 0.1** | **3.8 ± 0.1** |
| K17060 | Sensor regulator | two-component system, sensor histidine<br>kinase AauS [EC:2.7.13.3] | 76045 | 28 % | 70 % | **32 ± 5** | **45 ± 6** | 3.0 ± 0.1 | 3.0 ± 0.2 |
| K05876 | Sensor regulator | methyl-accepting chemotaxis protein III,<br>ribose and galactose sensor receptor | 38854 | 19 % | 79 % | 15 ± 2 | 24 ± 2 | **2.2 ± 0.1** | **2.5 ± 0.2** |
| K19611 | Sensor regulator | ferric enterobactin receptor | 3275 | 9 % | 91 % | **2 ± 2** | **4 ± 2** | 0.4 ± 0.4 | 0.6 ± 0.5 |
| K07786 | Sensor regulator | MFS transporter, DHA2 family, multidrug<br>resistance protein | 2193 | 15 % | 82 % | **3 ± 1** | **6 ± 1** | 1.0 ± 0.3 | 1.1 ± 0.4 |
| K06080 | Sensor regulator | RcsF protein | 572 | 5 % | 95 % | **0 ± 0** | **1 ± 0** | 0 ± 0 | 0.0 ± 0.1 |
| K10235 | Membrane transport | alpha-glucoside transport system ATP-binding protein | 117671 | 27 % | 71 % | **40 ± 6** | **55 ± 6** | **3.0 ± 0.1** | **3.2 ± 0.1** |
| K16088 | Membrane transport | outer-membrane receptor for ferric<br>coprogen and ferric-rhodotorulic acid | 107196 | 22 % | 77 % | **36 ± 12** | **64 ± 11** | 2.7 ± 0.4 | 3.0 ± 0.2 |
| K16210 | Membrane transport | oligogalacturonide transporter | 89287 | 28 % | 70 % | **30 ± 3** | **35 ± 5** | 2.5 ± 0.2 | 2.5 ± 0.2 |
| K02532 | Membrane transport | MFS transporter, OHS family, lactose<br>permease | 58021 | 19 % | 79 % | **17 ± 1** | **24 ± 3** | 2.3 ± 0.1 | 2.3 ± 0.2 |
| K16552 | Membrane transport | polysaccharide biosynthesis/export protein<br>ExoF | 34802 | 26 % | 72 % | **13 ± 4** | **20 ± 4** | **1.8 ± 0.3** | **2.2 ± 0.2** |
| K11734 | Membrane transport | aromatic amino acid transport protein AroP | 24606 | 26 % | 64 % | **22 ± 3** | **27 ± 2** | **2.6 ± 0.1** | **2.4 ± 0.2** |
| K08156 | Membrane transport | MFS transporter, DHA1 family, arabinose<br>polymer utilization protein | 19358 | 19 % | 79 % | **12 ± 3** | **19 ± 3** | 1.9 ± 0.3 | 1.7 ± 0.4 |
| K10094 | Membrane transport | nickel transport protein | 12104 | 21 % | 78 % | **10 ± 2** | **15 ± 2** | 2.0 ± 0.2 | 1.9 ± 0.1 |
| K11934 | Membrane transport | outer membrane protein X | 9090 | 23 % | 73 % | **9 ± 2** | **11 ± 1** | **1.9 ± 0.2** | **1.4 ± 0.2** |
| K16553 | Membrane transport | succinoglycan exporter | 8372 | 15 % | 84 % | **5 ± 2** | **9 ± 2** | 1.3 ± 0.2 | 1.5 ± 0.4 |
| K16348 | Membrane transport | entericidin B | 3282 | 22 % | 77 % | **5 ± 1** | **10 ± 3** | **1.5 ± 0.2** | **1.9 ± 0.3** |
| K10975 | Membrane transport | allantoin permease | 1617 | 11 % | 86 % | 1 ± 1 | 1 ± 1 | 0.2 ± 0.4 | 0.1 ± 0.1 |
| K16347 | Membrane transport | entericidin A | 1086 | 21 % | 77 % | 3 ± 2 | 5 ± 1 | 0.7 ± 0.5 | 1.1 ± 0.2 |
| K11743 | Membrane transport | spermidine export protein MdtJ | 699 | 18 % | 79 % | 1 ± 2 | 5 ± 0 | 0.3 ± 0.5 | 1.2 ± 0.2 |
| K16696 | Membrane transport | exopolysaccharide (amylovoran) exporter | 409 | 11 % | 87 % | 0 ± 1 | 2 ± 1 | 0.0 ± 0.1 | 0.3 ± 0.3 |
| K12285 | Biofilm formation | MSHA biogenesis protein MshO | 25345 | 27 % | 70 % | 17 ± 3 | 24 ± 4 | 2.5 ± 0.1 | 2.6 ± 0.1 |
| K12280 | Biofilm formation | MSHA biogenesis protein MshJ | 20898 | 26 % | 70 % | 14 ± 1 | 20 ± 4 | 2.3 ± 0.1 | 2.1 ± 0.1 |
| K12284 | Biofilm formation | MSHA biogenesis protein MshN | 17259 | 24 % | 71 % | 10 ± 2 | 14 ± 2 | 1.8 ± 0.1 | 1.6 ± 0.1 |
| K10927 | Biofilm formation | MSHA pilin protein MshD | 15335 | 27 % | 69 % | 13 ± 2 | 21 ± 2 | 2.2 ± 0.2 | 2.6 ± 0.1 |
| K20961 | Biofilm formation | diguanylate cyclase [EC:2.7.7.65] | 14724 | 21 % | 77 % | 9 ± 1 | 11 ± 2 | 1.4 ± 0.3 | 1.1 ± 0.2 |
| K10926 | Biofilm formation | MSHA pilin protein MshC | 11966 | 25 % | 71 % | 13 ± 2 | 18 ± 2 | 2.3 ± 0.2 | 2.4 ± 0.1 |
| K12286 | Biofilm formation | MSHA biogenesis protein MshP | 11500 | 26 % | 71 % | 10 ± 2 | 16 ± 2 | 2.0 ± 0.2 | 2.4 ± 0.1 |
| K12281 | Biofilm formation | MSHA biogenesis protein MshK | 4887 | 23 % | 74 % | 6 ± 1 | 9 ± 2 | 1.5 ± 0.1 | 1.3 ± 0.2 |
| K19731 | Quorum sensing | LuxR family transcriptional regulator, quorum-sensing system regulator CciR | 53445 | 27 % | 70 % | 37 ± 4 | 51 ± 7 | 3.0 ± 0.2 | 3.1 ± 0.1 |
| K20326 | Quorum sensing | protein XagA | 29432 | 23 % | 74 % | 17 ± 4 | 23 ± 3 | 2.0 ± 0.2 | 2.2 ± 0.2 |
| K20268 | Quorum sensing | rhizosphere induced protein | 10459 | 25 % | 72 % | 12 ± 4 | 23 ± 4 | 2.0 ± 0.4 | 2.4 ± 0.3 |
| K20267 | Quorum sensing | type IV secretion system protein TrbH | 3586 | 21 % | 78 % | 4 ± 1 | 6 ± 2 | 1.1 ± 0.3 | 1.5 ± 0.3 |
| K20275 | Quorum sensing | nematocidal protein AidA | 2480 | 22 % | 76 % | 4 ± 1 | 8 ± 2 | 1.1 ± 0.2 | 1.4 ± 0.2 |
| K20272 | Quorum sensing | TraR antiactivator | 1414 | 22 % | 78 % | 3 ± 1 | 5 ± 1 | 0.8 ± 0.4 | 1.4 ± 0.3 |
| K03477 | Transcriptional regulation | DeoR family transcriptional regulator, ulaG and ulaABCDEF operon transcriptional repressor | 93668 | 28 % | 68 % | 49 ± 5 | 55 ± 5 | 3.1 ± 0.1 | 2.9 ± 0.1 |
| K21825 | Transcriptional regulation | AraC family transcriptional regulator, L-arginine-responsive activator | 31901 | 25 % | 71 % | 26 ± 3 | 32 ± 2 | 2.7 ± 0.1 | 2.6 ± 0.2 |
| K05372 | Transcriptional regulation | AraC family transcriptional regulator | 5092 | 24 % | 74 % | 8 ± 3 | 14 ± 2 | 1.7 ± 0.4 | 2.1 ± 0.2 |
| K19060 | Transcriptional regulation | TetR/AcrR family transcriptional regulator, macrolide resistance operon repressor | 3198 | 22 % | 76 % | 6 ± 2 | 11 ± 2 | 1.5 ± 0.4 | 1.7 ± 0.2 |
| K12820 | Transcriptional regulation | pre-mRNA-splicing factor ATP-dependent RNA helicase DHX15/PRP43 [EC:3.6.4.13] | 1417 | 11 % | 87 % | **1 ± 1** | **2 ± 1** | 0.1 ± 0.3 | 0.2 ± 0.2 |
| K12059 | Secretion systems | conjugal transfer pilus assembly protein TrbC | 17109 | 26 % | 72 % | **18 ± 3** | **23 ± 6** | 2.4 ± 0.2 | 2.5 ± 0.3 |
| K12688 | Secretion systems | autotransporter serine protease [EC:3.4.21.- ] | 10969 | 18 % | 80 % | **7 ± 2** | **16 ± 3** | **1.5 ± 0.2** | **1.9 ± 0.4** |
| K11017 | Secretion systems | hemolysin activation/secretion protein | 9861 | 24 % | 73 % | **12 ± 3** | **17 ± 2** | 2.2 ± 0.1 | 2.0 ± 0.3 |
| K18380 | Secretion systems | type III secretion control protein HpaB | 5890 | 21 % | 77 % | **3 ± 1** | **7 ± 3** | **1.0 ± 0.2** | **1.3 ± 0.3** |
| K12069 | Secretion systems | conjugal transfer pilus assembly protein TraA | 5753 | 23 % | 75 % | 9 ± 3 | 11 ± 1 | 1.9 ± 0.2 | 1.7 ± 0.2 |
| K04338 | Secretion systems | curli production assembly/transport component CsgF | 4827 | 27 % | 68 % | **9 ± 2** | **16 ± 3** | **2.0 ± 0.2** | **2.3 ± 0.3** |
| K18373 | Secretion systems | type III secretion protein HrpB1 | 4535 | 23 % | 75 % | **5 ± 1** | **8 ± 2** | 1.3 ± 0.2 | 1.5 ± 0.3 |
| K11889 | Secretion systems | type VI secretion system protein ImpN [EC:2.7.11.1] | 4012 | 15 % | 85 % | **3 ± 2** | **7 ± 1** | **0.6 ± 0.5** | **1.3 ± 0.1** |
| K11909 | Secretion systems | type VI secretion system protein VasI | 2984 | 20 % | 79 % | **6 ± 3** | **10 ± 3** | 1.4 ± 0.4 | 1.6 ± 0.4 |
| K03202 | Secretion systems | type IV secretion system protein VirB7 | 2396 | 22 % | 77 % | **5 ± 2** | **8 ± 3** | 1.2 ± 0.5 | 1.6 ± 0.6 |
| K12083 | Secretion systems | type IV secretion system protein PtlH [EC:7.4.2.8] | 1536 | 20 % | 76 % | **3 ± 1** | **4 ± 0** | 0.8 ± 0.3 | 0.6 ± 0.2 |
| K12228 | Secretion systems | TrbB protein | 614 | 17 % | 82 % | **1 ± 1** | **4 ± 2** | **0.1 ± 0.2** | **1.2 ± 0.4** |
| K20555 | Secretion systems | type IV secretion system protein TrbK | 613 | 21 % | 79 % | **1 ± 1** | **3 ± 1** | **0.2 ± 0.3** | **0.9 ± 0.4** |
| K13454 | Secretion systems | type III effector protein AvrRpm1 | 77 | 0 % | 100 % | **0 ± 0** | **0 ± 1** | 0 ± 0 | 0 ± 0 |
| K19216 | Antibiotic resistance | metallo-beta-lactamase class B IND [EC:3.5.2.6] | 8968 | 18 % | 80 % | **4 ± 2** | **9 ± 2** | **0.7 ± 0.4** | **1.2 ± 0.4** |
| K18793 | Antibiotic resistance | beta-lactamase class D OXA-23 [EC:3.5.2.6] | 6092 | 9 % | 90 % | 3 ± 1 | 3 ± 1 | **0.5 ± 0.4** | **0.1 ± 0.0** |
| K19101 | Antibiotic resistance | beta-lactamase class C FOX [EC:3.5.2.6] | 5827 | 20 % | 73 % | **6 ± 2** | **9 ± 1** | **1.4 ± 0.3** | **1.7 ± 0.2** |
| K22335 | Antibiotic resistance | beta-lactamase class D OXA-114 [EC:3.5.2.6] | 4627 | 22 % | 76 % | **7 ± 3** | **13 ± 3** | **1.6 ± 0.5** | **2.0 ± 0.3** |
| K19213 | Antibiotic resistance | beta-lactamase class D OXA-12 [EC:3.5.2.6] | 1995 | 17 % | 80 % | **3 ± 1** | **8 ± 2** | **0.7 ± 0.5** | **1.6 ± 0.2** |
| K02399 | Motility | flagellar biosynthesis protein FlgN | 22038 | 24 % | 72 % | **19 ± 3** | **28 ± 5** | **2.4 ± 0.2** | **2.1 ± 0.2** |
| K07345 | Motility | major type 1 subunit fimbrin (pilin) | 9256 | 25 % | 72 % | **13 ± 4** | **23 ± 3** | 2.2 ± 0.3 | 2.5 ± 0.2 |
| K04643 | Secondary messaging | sensory rhodopsin | 6337 | 9 % | 89 % | **4 ± 2** | **8 ± 2** | 1.1 ± 0.5 | 1.0 ± 0.5 |
| K20966 | Secondary messaging | c-di-GMP phosphodiesterase [EC:3.1.4.52] | 4168 | 19 % | 79 % | **5 ± 1** | **7 ± 2** | **1.3 ± 0.2** | **1.6 ± 0.2** |
| K11964 | Defence | pellino [EC:2.3.2.27] | 2541 | 16 % | 84 % | **3 ± 2** | **6 ± 2** | **0.7 ± 0.6** | **1.5 ± 0.4** |
| K13964 | Defence | serpin B7 | 1767 | 16 % | 79 % | 3 ± 1 | 3 ± 1 | **0.5 ± 0.2** | **0.2 ± 0.2** |
| K08566 | Defence | plasminogen activator [EC:3.4.23.48] | 1624 | 17 % | 83 % | **2 ± 2** | **6 ± 1** | **0.5 ± 0.6** | **1.5 ± 0.2** |
| K01352 | Defence | granzyme A [EC:3.4.21.78] | 659 | 25 % | 72 % | **2 ± 1** | **5 ± 2** | **0.5 ± 0.5** | **1.3 ± 0.4** |
| K13448 | Calcium sensors | calcium-binding protein CML | 1065 | 16 % | 80 % | **1 ± 1** | **3 ± 1** | 0.2 ± 0.2 | 0.3 ± 0.2 |
| K10857 | Cell cycle | exodeoxyribonuclease X [EC:3.1.11.-] | 48331 | 27 % | 70 % | **26 ± 3** | **35 ± 5** | 2.1 ± 0.2 | 2.2 ± 0.2 |
| K13586 | Cell cycle | holdfast attachment protein HfaB | 29948 | 26 % | 72 % | 13 ± 1 | 14 ± 2 | **2.2 ± 0.1** | **2.0 ± 0.1** |
| K18642 | Cell cycle | crescentin | 10803 | 22 % | 78 % | **6 ± 1** | **7 ± 1** | 1.3 ± 0.3 | 1.2 ± 0.2 |
| K13585 | Cell cycle | holdfast attachment protein HfaA | 8477 | 19 % | 81 % | **7 ± 1** | **10 ± 1** | **1.7 ± 0.2** | **2.0 ± 0.2** |
| K14781 | Cell cycle | ATP-dependent RNA helicase DHR2 [EC:3.6.4.13] | 1029 | 8 % | 91 % | **0 ± 0** | **1 ± 0** | 0 ± 0 | 0 ± 0 |
| K13831 | Primary metabolism | 3-hexulose-6-phosphate synthase / 6-phospho-3-hexuloisomerase [EC:4.1.2.43 5.3.1.27] | 2008 | 56 % | 18 % | **3 ± 1** | **1 ± 1** | **1.0 ± 0.2** | **0.1 ± 0.3** |
| K00608 | Primary metabolism | aspartate carbamoyltransferase [EC:2.1.3.2] | 319 | 88 % | 10 % | **2 ± 1** | **0 ± 0** | **0.7 ± 0.5** | **0 ± 0** |
| K21346 | Primary metabolism | methionine transaminase [EC:2.6.1.88] | 311 | 62 % | 14 % | **2 ± 1** | **0 ± 0** | **0.5 ± 0.3** | **0 ± 0** |
| K05660 | Membrane transport | ATP-binding cassette, subfamily B (MDR/TAP), member 5 | 136 | 74 % | 7 % | 1 ± 1 | 0 ± 0 | 0.1 ± 0.2 | 0 ± 0 |
| K07991 | Motility | archaeal preflagellin peptidase FlaK [EC:3.4.23.52] | 602 | 22 % | 4 % | **1 ± 1** | **0 ± 0** | 0.2 ± 0.2 | 0 ± 0 |
| K02383 | Motility | flagellar protein FlbB | 55 | 97 % | 1 % | **0 ± 1** | **0 ± 0** | **0.1 ± 0.2** | **0 ± 0** |

The reads assigned to the differentially abundant functional genes between tall and semi-dwarf rhizospheres were re-processed though our bioinformatic pipeline and assigned to taxa. This allowed us to look for taxonomic differences and potential important taxa for these functions. Out of the 107 functional genes enriched in tall rhizospheres, 87 % had higher taxa richness in the tall rhizospheres, but only 37 % had higher diversity. This suggests that evenness in these communities may be important whereby dominant taxa could be driving these differences in function (Table 3). This was further evident by the differences in the community composition, based on the relative abundance of phyla, between tall and semi-dwarf rhizospheres for each differentially abundant gene (Table S2). In the tall rhizospheres the dominant phyla or class often contributed to a higher proportion of reads than in the semi-dwarf rhizospheres, except where a function was only carried out by taxa from a single phylum.

## Discussion

We compared the shotgun metagenomic profiles of prokaryote communities from the rhizospheres of tall and semi-dwarf wheat cultivars and found clear differences associated with wheat height phenotype. We also confirmed our hypothesis that taxonomic changes relate to changes in the potential function of these communities, with the main differences being driven by deterioration of many taxa and functional genes in semi-dwarf rhizospheres. Furthermore, the addition of unplanted bulk soil controls in our study confirm that the rhizosphere microbiome of semi-dwarf cultivars differentiates less from bulk soil than rhizosphere microbiomes derived from tall cultivars in terms of both taxonomy and function. These observations evidence that the incorporation of mutant *Rht* gene alleles which resulted in the semi-dwarf phenotype, have reduced the ability of these host plants to select a rhizosphere microbiome from the bulk soil microbial reservoir.

### 1) Taxonomic differences

When using shotgun metagenomics in contrast to phylogenetic marker gene analysis, we identified 10-fold more prokaryote taxa that were differentially abundant between the tall and semi-dwarf cultivars than in our previous study (Kavamura et al., 2020). Both studies found that most of the differentially abundant taxa belonged to Pseudomonadota, Bacteroidota and Actinomycetota and were enriched in the tall cultivars, but our results were to a much larger extent with 95 % enriched in tall compared with 69 % in Kavamura et al. (2020). We also found that species richness was higher in the semi-dwarf rhizospheres, supporting the conclusion that they are less selective than tall wheat. However, species diversity was higher in tall rhizospheres in our study and not in Kavamura et al. (2020). While the overall trend was the same in our previous work these minor discrepancies could also be the result of the different edaphic factors and different growing conditions.

We demonstrated a loss of Bacteroidota through wheat dwarfing both in relative abundance and in differentially abundant taxa. This has been found in wheat domestication from other studies (Aleklett et al., 2015; Pérez-Jaramillo et al., 2018; Rossmann et al., 2020) and Bacteroidota have been shown to be an important phylum for plant pathogen protection and phosphorus uptake (Lidbury et al., 2021). Domestication has also been shown to enrich Actinobacteria (Aleklett et al., 2015; Pérez-Jaramillo et al., 2018; Rossmann et al., 2020) and phyla that are more commonly associated with bulk soil, such as Acidobacteriota and Verrucomicrobiota (Kavamura et al., 2020). Our results showed a similar trend with Acidobacteriota and Verrucomicrobiota having a reduced relative abundance in semi-dwarf and tall rhizospheres compared with bulk soil, (Supplementary Figure 2) but there were taxa from these phyla enriched in the rhizospheres of tall cultivars and not in semi-dwarf rhizosphere samples. However, by including unplanted bulk soil controls our study confirms that the rhizosphere microbiomes of semi-dwarf wheat cultivars are more similar to bulk soils than the rhizosphere microbiomes of tall cultivars which would indicate less filtering of Prokaryote taxa in semi-dwarf wheat varieties.

### 2) Functional gene differences Bulk soil vs rhizosphere soils

As described in the results section, 19 of the 20 most abundant genes that are enriched in tall rhizospheres compared to bulk soils are also more abundant in tall rhizospheres compared to semi-dwarfs, but to a lesser extent (Table S1). This suggests that the selective ability of semi-dwarf plants is diminished in comparison to tall wheats, and the rhizosphere microbiomes in semi-dwarf are a ‘middle ground’ between bulk soil and tall rhizospheres (Fig. 1D). The one exception being K07305 (peptide-methionine (R)-S-oxide reductase) which is equally abundant in the rhizosphere of tall and short microbiomes and much higher compared to bulk soil samples (approximate mean normalised read count of 500K per sample for each height class and 48K for bulk soil). This enzyme plays a crucial role in repairing oxidatively damaged proteins. We hypothesise that its high abundance in the rhizosphere reflects the high metabolic activity of the rhizosphere with consequential generation of free radicals. It implies that the high incidence of this gene and presumably expression of the enzyme it encodes is crucial for rhizosphere competence regardless of wheat genotype. Future studies could also determine whether expression of this gene is imperative for microbial colonisation of the plant-root environment.

### Tall vs semi-dwarf rhizospheres

Shotgun metagenomic analysis of samples from the rhizosphere of tall and semi-dwarf wheat cultivars resulted in the detection of clear differential abundance in reads mapping to 113 genes. With 107 genes enriched in the rhizospheres of tall wheats, these results possibly indicate that the genetic potential of the host plant to influence the root microbiome structure and function has been reduced as a consequence of wheat dwarfing. We discuss the functions associated with genes enriched in the rhizospheres of tall wheats (see KEGG IDs in brackets) under 11 major categories below in order of highest read counts allocated per category. Not all 107 genes are discussed in detail but see Table 3 and Table S2 for a summary in terms of read counts and taxonomic information. The six genes enriched in the semi-dwarfs assign to primary metabolism, membrane transport and motility, but in much fewer read counts (3K reads) than those enriched in tall cultivars (6.5M reads).

#### Secondary metabolism

There were 13 differentially abundant genes relating to secondary metabolism with a total of 2.6M reads assigned over all samples. Of these reads, 98% were most similar to genes associated with biosynthesis of staurosporine (K14266, K19885, K20086, K20090 and K20088), a natural product antibiotic originally isolated from the bacterium *Streptomyces staurosporeus* (Omura et al., 1977), with mode of action being through competitive protein kinase inhibition, with this family of molecules exhibiting anti-cancer potential (Yadav et al., 2015). In addition, reads were associated with terpenoid and polyketide antibiotic synthesis (K16397, K16403, K14627, K14368, K21212, K16448 and K15968) as well as bacterial toxin production (K11009). The high number of differentially abundant genes associated with antibiotic production could indicate that niche occupancy competition of rhizosphere microbiome community members in pre-green revolution wheat is driven by an arms race, which could provide a novel underexploited resource for natural product discovery and the development of the next generation of antibiotics.

#### Cell wall degradation

A total of 1.3M reads were found to map to four differentially abundant genes for plant cell wall degradation (K18651, K18578, K18576 and K18786). Although this function has long been associated with plant pathology (Kubicek et al., 2014; Wood, 1960), cellulolytic activity has also been linked to enhanced plant root length by facilitating sloughing-off of root cap cells from the root tip which assists the growing root in penetrating soil (Campillo et al., 2004). It has also been found that cell wall degradation is essential for *Rhizobium* symbiotic infection of legume roots (Robledo et al., 2008). It follows that this function could also be important for microbial colonisation of the plant environment, microbial mediation of plant root architecture and an overwintering energy source for microbes on crop residues. With such high enrichment of these genes in tall wheat rhizospheres therefore demonstrates that a potentially important function to facilitate plant-microbial interactions has been degraded in semi-dwarf wheat.

#### Primary metabolism

In addition to cell wall degradation, a further 809K reads mapped to genes associated with primary metabolism. These can primarily be subdivided into the metabolism of carbohydrates (380K reads), lipids (340K reads), proteins (46K reads), co-factors/vitamins (27K reads), and enzymes involved in core primary metabolism (15K reads). These observations are congruent with the classes of genes most abundant in membrane transport, and it is unsurprising that genes associated with the metabolism of these primary nutrient sources are abundant in these data; however, it is interesting that they are highly enriched in tall rhizospheres. The most abundant of the differentially abundant primary metabolism genes was associated with sphingolipid metabolism through the action of galactosylceramidase (K01202, 336K reads). Sphingolipids are a class of membrane bound lipid which act as signal bioactive molecules (Wang et al., 2021b). It has been proposed that Bacteroidota, predominant members of the mammalian gut microbiome, utilise sphingolipids as an energy source as well as to mediate signal transduction and stress response pathways, facilitating their persistence in this environment (An et al., 2014). Our dataset found that Actinomycetota made up the highest proportion of taxa with this gene (47%) demonstrating it is not a Bacteroidota specific function in the wheat root environment. It could be the case that microbe-microbe and microbe-plant sphingolipid-based signalling is also crucial in commensal colonisation of the plant root environment as is proposed in the human gut. There were also two genes involved in propanoate metabolism (K20455, K08325) which is a building block of fatty acids which are often a component of sphingolipids, the genes for their biosynthesis are highly enriched in the tall rhizospheres in this dataset and have previously been found to be enriched in the metabolism pathways of rhizosphere microbiota (Finkel et al., 2017; Pascale et al., 2020; Zhou et al., 2020).

Furthermore, one of the differentially abundant carbohydrate metabolism genes was associated with lipopolysaccharide (LPS) biosynthesis (K02847). LPS is an important outer membrane component of Gram-negative bacteria and mutations in LPS biosynthetic genes have been shown to impair rhizosphere and root colonisation of maize by *Rhizobium tropici,* a root nodule bacterium (Ormeño-Orrillo et al., 2012). Finally, we also found K08961, which encodes a gene for glycosaminoglycan degradation to differentially more abundant in tall wheat rhizosphere microbiomes. This linear polysaccharide is not present in plant cells, however it is found in animal as well as bacterial cells, so its function in the root environment is likely to be implicated in nutrient cycling of microbiome community members.

#### Sensor regulator

Approximately 640K reads were found to map to differentially abundant genes associated with two component sensor-regulation systems, most of which (510K reads) were assigned to serine detection (K05874). In addition, histidine kinase AauS (K17060), which regulates uptake and metabolism of acidic amino acids in *Pseudomonas putida* (Sonawane et al., 2006) in a phosphorylation dependent manner, was also in high abundance, along with a sensor receptor gene for monosaccharides (ribose and galactose; K05876). It will be interesting in future work to determine the amino acid to sugar ratio and the relative contribution of amino acids in the root exudates of tall compared to dwarfed wheats and determine whether detection of these molecules is important for chemoattraction into the root zone as a prerequisite to rhizosphere colonisation of tall wheat.

#### Membrane transport

A total of 520K reads mapped to differentially abundant genes associated with membrane transport and the vast majority (435K reads) were associated with uptake systems. However, export systems were largely associated with polysaccharide transport, presumably as a prerequisite to biofilm formation (e.g. K16552, K16553, K16696), though the outer membrane protein X (K11934), the entericidin toxin peptide and antidote system (K16347, K16348) and spermidine export system (K11743; which is important at acidic pH such as that of the rhizosphere (Higashi et al., 2008) are also represented). Regarding import systems, these were categorised as metal (iron and nickel; K10094, K16088), purine (K10975), aromatic amino acids (K11734), though the vast majority of these reads were associated with ATP-binding cassette and MFS sugar uptake systems (alpha-glycosides, arabinose, lactose and oligogalacturonides) with a total of 280K reads assigned (K02532, K10235, K16210, K08156). The observation that bacteria in the rhizosphere have a high level of uptake transport systems has previously been studied in rhizobia (Mauchline et al., 2006), though this is the first time that an enhancement in these systems is associated with the root microbiome of pre-green revolution wheats. It is interesting that there was no perceived difference in abundance detected between tall and short wheats for genes associated with organic acid uptake systems and implies that the dwarfing of wheat has perhaps not impacted the root exudation profile of these molecules to the same extent of amino acids and sugars.

It was also found that the ratio of reads associated with amino acid:sugar detection in two component systems is inversely proportional to the ratio of predicted uptake transport systems for these substrates. It is possible that as sugars and organic acids are more ubiquitous in root exudates, that the root microbiome of tall wheats have evolved to invest a greater proportion of detection mechanisms into amino acid detection. Future work should determine the differential root exudate composition between tall and semi-dwarf wheats and whether these can be positively correlated with the predicted uptake systems.

#### Biofilm formation and quorum sensing

Approximately 234K reads mapped to genes associated with quorum sensing and biofilm formation. Reads mapping to genes associated with mannose sensitive haemagglutinin (MSHA) pilus biogenesis [*mshJ*, (K12280) *mshK* (K12281), *mshN* (K12284), *mshO* (K12285) and *mshP* (K12286)] are total 83K reads. This molecule has been shown to be crucial for the attachment of *Vibrio cholerae* to environmental surfaces (Marsh and Taylor, 1999) and more recently MSHA has also been shown to promote the attachment of *Pseudoalteromonas tunicata* to cellulose surfaces (Dalisay et al., 2006). In addition, approximately 28K reads are assigned to MSHA pilin protein production genes *mshC* and *mshD* (K10926, K10927). These genes are important for biofilm formation, horizontal gene transfer and twitching motility (Toma et al., 2002). It therefore follows that the increased differential abundance of MSHA could reflect their importance for microbial persistence in the tall wheat rhizosphere microbiome.

In addition, approximately 106K reads mapped to genes associated with quorum sensing. The most abundant of these, at 56K reads, being assigned as the LuxR family transcriptional regulator gene *cciR* (K19731). A study into the food-borne pathogen *Vibrio parhaemolyticus* revealed that LuxR family regulators influence bacterial growth and biofilm formation and highlighted that the LuxR regulator RobA regulated expression of exopolysaccharides (EPS) synthesis and controlled biofilm formation (Zhong et al., 2021). Our data suggests that transcriptional regulators are also important for biofilm formation in the root environment and that the increased differential abundance in the rhizosphere of tall cultivars is indicative of a tighter control of rhizosphere microbiome assembly in pre green revolution wheat.

#### Transcriptional regulation

Approximately 142K reads were assigned to transcriptional regulation, and almost 100K are ascribed to the DeoR family transcriptional regulator (K03477). In *E. coli*, a DeoR family transcriptional regulator, UlaR, was found to be responsible for suppressing transcription of the divergent *ulaG* and *ulaABCDEF* operons (which catabolise L-ascorbate), under ascorbate depleted conditions (Campos et al., 2004). Interestingly, ascorbate has been shown to be released from plant roots under conditions of salt stress and influences root elongation (Li et al., 2018; Makavitskaya et al., 2018). It could follow that exudation of ascorbate into the rhizosphere profoundly affects root microbiome colonisation patterns that are employed by tall wheat cultivars. In addition, 33K reads were assigned to the AraC L-arginine responsive activator (K21825). It has previously been shown that arginine is an important environmental cue for cyclic di-GMP signalling (c-di-GMP) signalling and biofilm formation in *Pseudomonas putida* (Barrientos-Moreno et al., 2020), highlighting the potential importance of detection and utilisation of this amino acid for microbiome assembly in the rhizosphere of tall wheat. It will be interesting to assess whether tall cultivars produce a greater proportion of arginine in their exudates than the corresponding short cultivars.

#### Secretion systems

A total of 76K reads mapped to secretion systems, and interestingly, type III (K13454, K18380, K18373, K18380), IV (K03202, K12083, K120555) and type VI (K11889, K11909) secretion systems are predominately over represented in tall wheat rhizosphere, all of which use machinery to directly breach and deliver secreted proteins across host cell membranes as opposed to other secretion system types which release toxins into the extracellular milieu (Green and Mecsas, 2016). In addition, genes associated with pilus assembly proteins for conjugal transfer (a total of 25K reads; K12059, K12069) were also enriched in tall rhizospheres and are key to bacterial mating via synthesis of a type IV secretion system (Shala-Lawrence et al., 2018).

#### Antibiotic resistance

Approximately 29K reads were assigned to differentially abundant genes involved in antibiotic resistance (K19216, K18793, K19101, K22335 & K19213). These were related to beta-lactamase function, which has been shown to be well represented in isolates of the soil dwelling bacterium *Bacillus subtilis* (Bucher et al., 2019). Evidence suggests that its production facilitates rhizosphere colonisation by the plant pathogen *Fusarium oxysporum* (Chang et al., 2021). It is interesting that the number of reads in this category is far exceeded by the number of reads for antibiotic production (e.g. staurosporine biosynthesis). This could indicate that microbial strategy for rhizosphere colonisation of tall wheat rhizospheres has an offensive emphasis, and that some of this capacity is reduced in the short wheat rhizosphere microbiome.

#### Motility

There were two differentially abundant genes relating to motility, but the majority of the reads (23K reads) were assigned to the flagellar chaperone biosynthesis gene *flgN* (K02399). This gene has been shown to be required for flagellum-based motility in *Bacillus subtilis* (Cairns et al., 2014) and is involved in the regulation and assembly of the flagellum, and its enhanced differential abundance in the rhizosphere of tall wheat suggests that it is important for the colonisation of this environment.

#### Secondary messaging

Approximately 10K reads were assigned to secondary messaging, especially cyclic di GMP (c-di-GMP) phosphodiesterase (K20966) which has previously been shown to be important for the rhizosphere colonisation of wheat by *Pseudomonas fluorescens* (Little et al., 2019).

The activity of c-di-GMP is important for the production of EPS and biofilm formation and an increase in abundance of c-di-GMP phosphodiesterase activity which is involved in the reduction of c-di-GMP level implies that biofilm formation in tall cultivars is tightly regulated.

### Conclusions

Our data showed that semi-dwarf wheat has a reduced rhizosphere effect when compared with tall wheat. In addition, we observed a depletion of a wide range of functional genes in semi-dwarf wheat, indicating a functional deterioration in the rhizosphere microbiome associated with wheat dwarfing. Wheat dwarfing of the green revolution has profoundly influenced the selection and function of the root microbiome, and this is evidenced by reduced abundance in genes such as those involved in biofilm formation as well as the sensing of key nutrients and their transport into microbial cells in semi-dwarf wheat.

However, the vast majority of reads map to genes associated with secondary metabolism as well as cell wall degradation. As such it seems that the root microbiome of tall wheats have adopted a two-pronged strategy of exclusion of microbial competitor reduction through antibiotic production (e.g. staurosporine production) as well as the metabolism of plant derived nutrients. The latter seems to be via utilisation of cell wall constituents as a nutrient source to establish in this environment as well as through sphingolipid metabolism – it is unclear to what extent these functions are also important for plant-microbe signalling. The green revolution combined wheat dwarfing with the application of synthetic chemical fertilisers. Our previous work has highlighted that application of synthetic fertiliser reduces the selection of nutrient-solubilising bacteria in the rhizosphere (Reid et al., 2021). It will be interesting to ascertain the impact of the combination of these factors for global microbiome function and selection, and to determine the specific contribution of *Rht* mutant alleles in isogenic wheats to the observed microbiome shifts caused by dwarfing which are highlighted in this work. Finally, we believe our results are of striking importance and highlight that implementation of microbiome facilitated agriculture as part of a sustainable crop production strategy will require an overhaul of wheat breeding programmes to consider plant-microbe interactions, especially in the root environment.

## Declarations

### Ethics approval and consent to participate

Not applicable

### Consent for publication

Not applicable

### Availability of data and material

The datasets generated and analysed during the current study are available in the NCBI repository, BioProject PRJNA1128034. The BioProject and associated SRA metadata are publicly available at https://www.ncbi.nlm.nih.gov/bioproject/PRJNA1128034. However, due to a glitch in the SRA system the samples are only available via individual download until this issue is resolved, see links below.

## BioSample

SAMN42186182 DNA_from_cv.Chidham_rep1 - BioSample - NCBI

SAMN42186183 DNA_from_cv.Chidham_rep2 - BioSample - NCBI

SAMN42186184 DNA_from_cv.Chidham_rep3 - BioSample - NCBI

SAMN42186185 DNA_from_cv.Chidham_rep4 - BioSample - NCBI

SAMN42186186 DNA_from_cv.Chidham_rep5 - BioSample - NCBI

SAMN42186187 DNA_from_cv.Gallant_rep1 - BioSample - NCBI

SAMN42186188 DNA_from_cv.Gallant_rep2 - BioSample - NCBI

SAMN42186189 DNA_from_cv.Gallant_rep3 - BioSample - NCBI

SAMN42186190 DNA_from_cv.Gallant_rep4 - BioSample - NCBI

SAMN42186191 DNA_from_cv.Gallant_rep5 - BioSample - NCBI

SAMN42186192 DNA_from_cv.Malacca_rep1 - BioSample - NCBI

SAMN42186193 DNA_from_cv.Malacca_rep2 - BioSample - NCBI

SAMN42186194 DNA_from_cv.Malacca_rep3 - BioSample - NCBI

SAMN42186195 DNA_from_cv.Malacca_rep4 - BioSample - NCBI

SAMN42186196 DNA_from_cv.Malacca_rep5 - BioSample - NCBI

SAMN42186197 DNA_from_cv.Red_Lammas_rep1 - BioSample - NCBI

SAMN42186198 DNA_from_cv.Red_Lammas_rep2 - BioSample - NCBI

SAMN42186199 DNA_from_cv.Red_Lammas_rep3 - BioSample - NCBI

SAMN42186200 DNA_from_cv.Red_Lammas_rep4 - BioSample - NCBI

SAMN42186201 DNA_from_cv.Red_Lammas_rep4 - BioSample - NCBI

SAMN42186202 DNA_from_Bulk_Soil_rep1 - BioSample - NCBI

SAMN42186203 DNA_from_Bulk_Soil_rep2 - BioSample - NCBI

SAMN42186204 DNA_from_Bulk_Soil_rep3 - BioSample - NCBI

**SRA**

SRR29660204 DNA_from_644 cv.Chidham_rep1 - SRA - NCBI

SRR29660203 DNA_from_cv.Chidham_rep2 - SRA - NCBI

SRR29660192 DNA_from_cv.Chidham_rep3 - SRA - NCBI

SRR29660188 DNA_from_cv.Chidham_rep4 - SRA - NCBI

SRR29660187 DNA_from_cv.Chidham_rep5 - SRA - NCBI

SRR29660186 DNA_from_cv.Gallant_rep1 - SRA - NCBI

SRR29660185 DNA_from_cv.Gallant_rep2 - SRA - NCBI

SRR29660184 DNA_from_cv.Gallant_rep3 - SRA - NCBI

SRR29660183 DNA_from_cv.Gallant_rep4 - SRA - NCBI

SRR29660182 DNA_from_cv.Gallant_rep5 - SRA - NCBI

SRR29660202 DNA_from_cv.Malacca_rep1 - SRA - NCBI

SRR29660201 DNA_from_cv.Malacca_rep2 - SRA - NCBI

SRR29660200 DNA_from_cv.Malacca_rep3 - SRA - NCBI

SRR29660199 DNA_from_cv.Malacca_rep4 - SRA - NCBI

SRR29660198 DNA_from_cv.Malacca_rep5 - SRA - NCBI

SRR29660197 DNA_from_cv.Red_Lammas_rep1 - SRA - NCBI

SRR29660196 DNA_from_cv.Red_Lammas_rep2 - SRA - NCBI

SRR29660195 DNA_from_cv.Red_Lammas_rep3 - SRA - NCBI

SRR29660194 DNA_from_cv.Red_662 Lammas_rep4 - SRA - NCBI

SRR29660193 DNA_from_cv.Red_Lammas_rep5 - SRA - NCBI

SRR29660191 DNA_from_Bulk_Soil_rep1 - SRA - NCBI

SRR29660190 DNA_from_Bulk_Soil_rep2 - SRA - NCBI

SRR29660189 DNA_from_Bulk_Soil_rep3 - SRA - NCBI

## Authors’ contributions

T.H.M., R.M. and M.E.S. acquired funding. T.H.M, I.M.C. and V.N.K. designed the experiments and collected the data. Bioinformatic analysis was performed by D.H. and M.E.S performed statistical analysis. Results were interpreted by T.H.M, I.M.C, G.L. and M.E.S. The manuscript was written by M.E.S. and T.H.M. All co-authors edited and commented on the manuscript.

## Acknowledgements

The authors have no further acknowledgements to include.

