## Appendix 1 for "Uncovering functional deterioration in the rhizosphere microbiome associated with wheat dwarfing"

### CONTENTS

### SI 1. SUPPLEMENTARY FIGURES

**Fig. S1.** Rarefaction curves for all samples based on the a) taxonomy and function b) of the prokaryote communities.

**Fig. S2.** Relative abundance of Prokaryote taxa in each metagenomic sample (S1-S23).

**Fig. S3.** The proportion (%) of enriched taxa belonging to each phylum according to differential abundance analysis contrasts between each pair of sample types.

### SI 2. SUPPLEMENTARY TABLES

**Table S1.** The 20 most abundant functional genes out of 1,712 that are differentially abundant between bulk soils and tall wheat rhizospheres.

**Table S2.** Mean relative abundance (%) of reads assigned to the most common phyla for the differentially abundant functional genes between tall and semi-dwarf rhizospheres.

### SI 1. SUPPLEMENTARY FIGURES

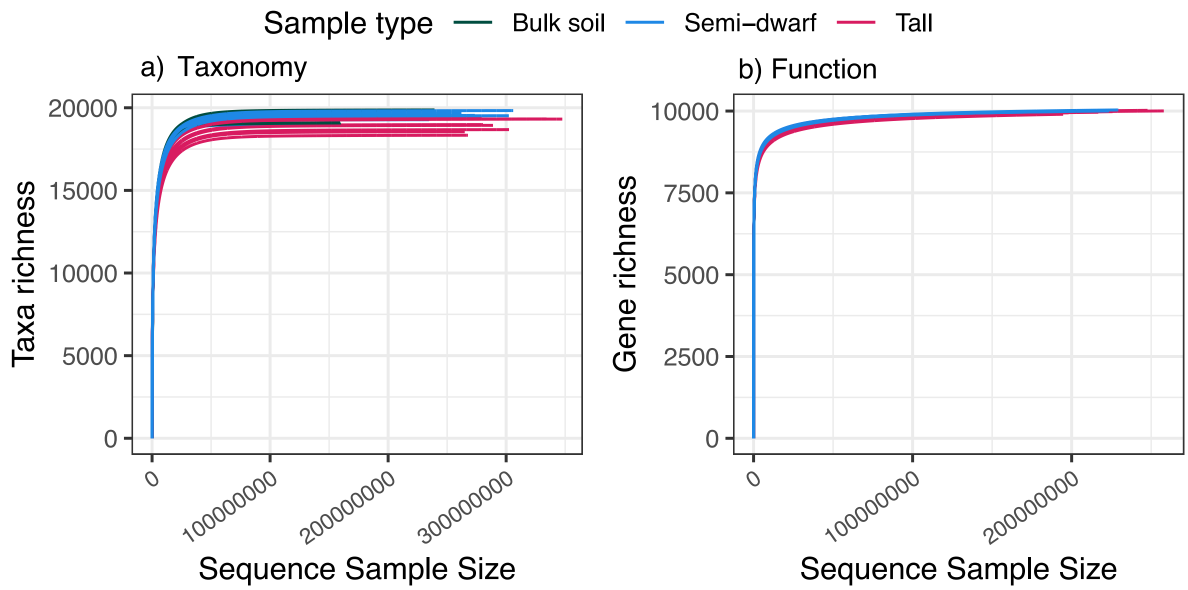


**Fig. S1. Rarefaction curves for all samples based on the a) taxonomy and function b) of the prokaryote communities.** Sample types, i.e., bulk soil controls or rhizosphere soil from semi-dwarf or tall wheat cultivars, are depicted by the colours of the curves.

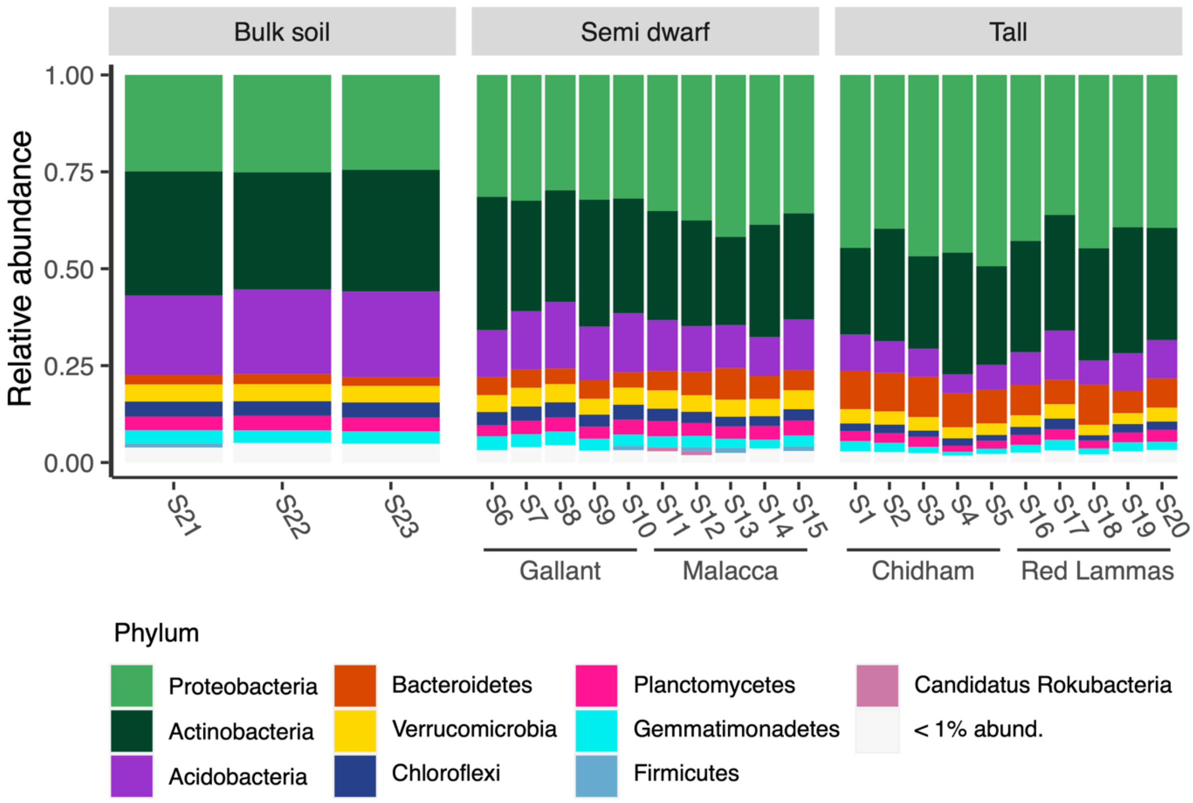


**Fig. S2. Relative abundance of Prokaryote taxa in each metagenomic sample (S1-S23)**. Colours represent the phyla level of the most abundant (> 1 % relative abundance) and taxa not classified at this level were excluded from analysis. Samples are grouped first by sample type, (grey header boxes) and then by wheat cultivar (x-axis labels).

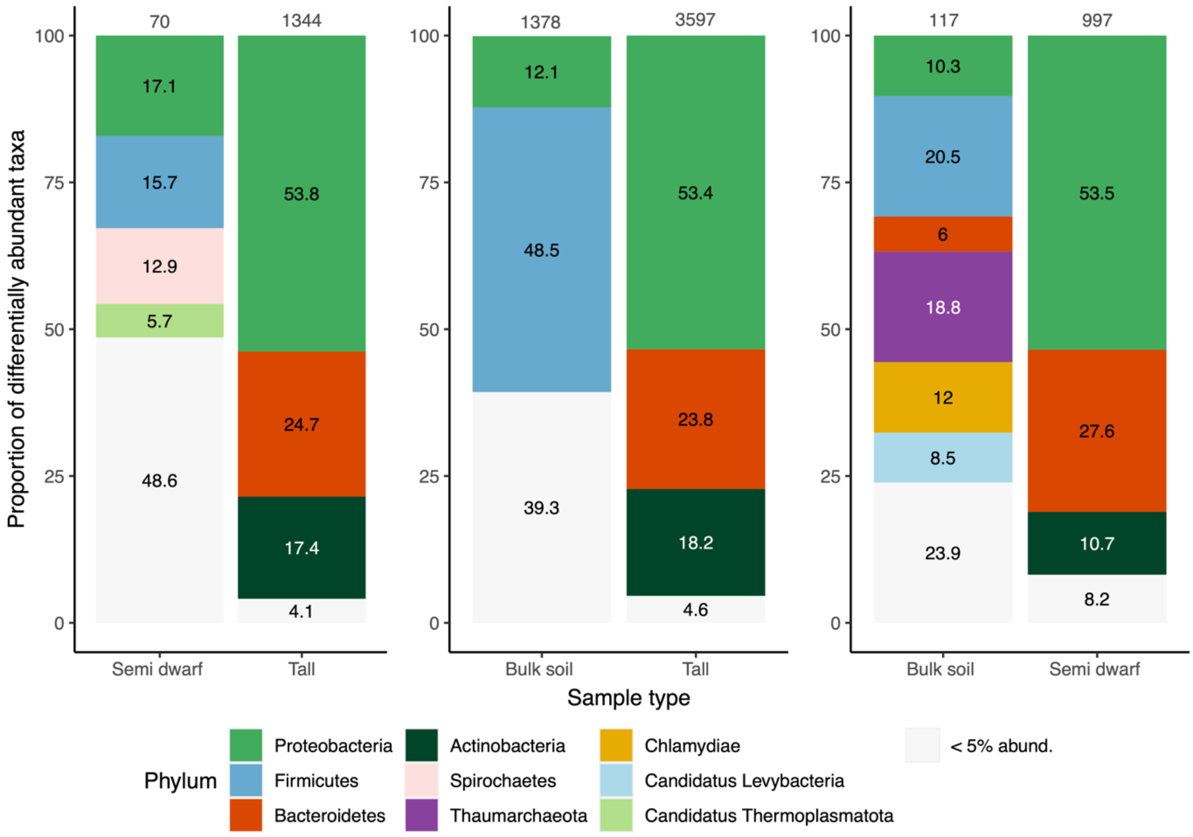


**Fig. S3. The proportion (%) of enriched taxa belonging to each phylum according to differential abundance analysis contrasts between each pair of sample types.** The proportions were based on the total number of taxa enriched in each sample type per comparison (numbers on top of bars) and each plot represents a different comparison with the sample types on the x-axis. Only phyla that contained more than 5% of taxa are indicated by colour, all others were considered rare and grouped together (< 5% abund.).

### SI2. SUPPLEMENTARY TABLES

**Table S1. The 20 most abundant functional genes out of 1,712 that are differentially abundant between bulk soils and tall wheat rhizospheres.** The functional genes are identified by the Kyoto Encyclopaedia of Genes and Genomes identifier (KEGG ID) and put into broad functional groups, and secondary functions are included where applicable. The total read count refers to the total number of reads for each functional gene found in rhizosphere samples normalised by DESeq2 size factors for each sample (McMurdie and Holmes, 2013). The means for each sample type (bulk soil and rhizosphere soil from semi-dwarf or tall wheat) were compared and significant differences are depicted by different letters (ANOVA, $p < 0.05$).

| KEGG ID | Functional group | Secondary function | Gene | Total read count (normalised) | Bulk soil mean (n=3) | Semi-dwarf mean (n=10) | Tall mean (n=10) |
|---|---|---|---|---|---|---|---|
| K02014 | Membrane transport | Iron transport | iron complex outermembrane recepter protein | 24,116,914 | 410,340a | 774,734b | 1,408,208c |
| K07305 | Primary metabolism | | peptide-methionine (R)-S-oxide reductase [EC:1.8.4.12] | 10,282,206 | 48,158a | 500,964b | 498,096b |
| K21573 | Membrane transport | | TonB-dependent starch-binding outer membrane protein SusC | 7,793,265 | 99,172a | 245,353a | 472,111b |
| K02529 | Transcription factors | | LacI family transcriptional regulator | 7,145,118 | 158,668a | 253,414b | 388,382c |
| K03406 | Two-component system | | methyl-accepting chemotaxis protein | 6,392,258 | 115,927a | 212,267a | 367,048b |

| | | | | | | | |
|---|---|---|---|---|---|---|---|
| K01190 | Primary metabolism | Carbohydrate metabolism | beta-galactosidase [EC:3.2.1.23] | 5,975,470 | 127,700a | 196,772a | 341,105b |
| K21572 | Membrane transport | | starch-binding outer membrane protein, SusD/RagB family | 5,364,121 | 79,015a | 170,800a | 320,784b |
| K03566 | Biofilm formation | | LysR family transcriptional regulator, glycine cleavage system transcriptional activator | 2,563,859 | 44,755a | 88,709b | 144,466c |
| K12308 | Primary metabolism | Carbohydrate metabolism | beta-galactosidase [EC:3.2.1.23] | 2,560,477 | 53,670a | 84,359a | 146,379b |
| K14266 | Secondary metabolism | Staurosporine biosynthesis | tryptophan 7-halogenase [EC:1.14.19.9] | 2,558,916 | 19,996a | 65,155b | 170,581c |
| K01811 | Primary metabolism | Carbohydrate metabolism | alpha-D-xyloside xylohydrolase [EC:3.2.1.177] | 2,386,478 | 57,423a | 81,787a | 131,742b |
| K01198 | Primary metabolism | Carbohydrate metabolism | xylan 1,4-beta-xylosidase [EC:3.2.1.37] | 2,361,118 | 45,804a | 79,884b | 133,775c |
| K00362 | Primary metabolism | Nitrogen metabolism | nitrite reductase (NADH) large subunit [EC:1.7.1.15] | 2,327,961 | 55,160a | 88,952b | 120,169c |
| K18900 | Transcription factors | | LysR family transcriptional regulator, regulator for bpeEF and oprC | 2,191,402 | 32,775a | 72,464b | 128,082c |
| K15923 | Primary metabolism | Carbohydrate metabolism | alpha-L-fucosidase 2 [EC:3.2.1.51] | 1,917,215 | 36,859a | 64,578b | 108,985c |
| K07552 | Membrane transport | | MFS transporter, DHA1 family, multidrug resistance protein | 1,860,700 | 44,058a | 70,312b | 96,325c |
| K12373 | Primary metabolism | Carbohydrate metabolism | hexosaminidase [EC:3.2.1.52] | 1,739,338 | 43,712a | 56,293a | 98,598b |
| K15270 | Membrane transport | | S-adenosylmethionine uptake transporter | 1,493,249 | 32,067a | 53,982b | 80,286c |
| K05970 | Primary metabolism | | sialate O-acetylesterase [EC:3.1.1.53] | 1,414,362 | 33,542a | 52,449b | 74,192c |
| K16135 | Transcription factors | | LysR family transcriptional regulator, transcriptional activator for dmlA | 1,377,784 | 19,342a | 44,193b | 81,970c |

**Table S2. Mean relative abundance (%) of reads assigned to the most common phyla for the differentially abundant functional genes between tall and semi-dwarf rhizospheres.** The relative abundances of Proteobacteria are shown at the class level whereby the three most dominant classes, alphaproteobacteria (α), betaproteobacteria (β) and gammaproteobacterial (γ) are shown and other classes are grouped together. Means were calculated for tall and semi-dwarf rhizosphere samples (n = 10). Dominant taxa group in each wheat height phenotype are underlined for each functional gene. Genes enriched in the tall and semi-dwarf rhizospheres are separated by the horizontal line. Results from PERMANOVAs based on Bray-Curtis dissimilarity of the relative abundances of all phyla and with wheat height phenotype as the main effect are presented. Significant differences between semi-dwarf and tall gene-associated communities are in bold and *p* values have been adjusted for multiple comparisons using a false-discovery rate correction. NA are present when there was not enough variation, i.e., the functional gene is only present in one phylum or if there were not enough samples in a particular height phenotype to run the model. See Table 3 for more details on each gene.

| | | **Semi-dwarf** | | | | | | | | | **Tall** | | | | | | | | | Permanova |
|---|---|---|---|---|---|---|---|---|---|---|---|---|---|---|---|---|---|---|---|---|
| | | Pseudomonadota | | | | Actino-mycetota | Bacter-iodota | Bacillota | Verruco-microbiota | Other | Pseudomonadota | | | | Actino-mycetota | Bacter-iodota | Bacillota | Verruco-microbiota | Other | (*p*-values) |
| KEGG ID | Function | α | β | γ | other | | | | | | α | β | γ | other | | | | | | |
| K14266 | Secondary metabolism | <u>58</u> | 14 | 6 | 14 | | | | 4 | 4 | <u>58</u> | 24 | 5 | 10 | | | | 2 | 1 | **<0.001** |
| K16397 | Secondary metabolism | | | | 2 | <u>80</u> | | | | 18 | | | | <1 | <u>94</u> | | | | 6 | **<0.001** |
| K16403 | Secondary metabolism | | | | <1 | <u>83</u> | | | | 16 | | | | | <u>96</u> | | | | 4 | **<0.001** |
| K14627 | Secondary metabolism | | | | | <u>100</u> | | | | | | | | | <u>100</u> | | | | | NA |
| K14368 | Secondary metabolism | | | | | <u>100</u> | | | | <1 | | | | | <u>100</u> | | | | <1 | **0.017** |

| | | | | | | | | | | | | | | | | | | | | |
|---|---|---|---|---|---|---|---|---|---|---|---|---|---|---|---|---|---|---|---|---|
| K11009 | Secondary metabolism | | | | 36 | 47 | | | | 17 | | 4 | | 12 | 83 | | | | 2 | **<0.001** |
| K19885 | Secondary metabolism | | 6 | | <1 | 58 | | | | 36 | | 24 | | <1 | 76 | | | | <1 | 0.050 |
| K21212 | Secondary metabolism | | | | | 99 | | | | <1 | | | | | 99 | | | | <1 | 0.593 |
| K16448 | Secondary metabolism | | | | | 51 | | | | 49 | | | | | 94 | | | | 6 | **<0.001** |
| K15968 | Secondary metabolism | | | | | 90 | | | | 10 | | | | | 100 | | | | | 0.288 |
| K20086 | Secondary metabolism | | | | | | | | | 100 | | 99 | | <1 | | | | | <1 | 1.000 |
| K20090 | Secondary metabolism | | | | | | | | | 100 | | 80 | | | | | | | 20 | **0.010** |
| K20088 | Secondary metabolism | | | | | | | | | 100 | | 69 | | 1 | | | | | 30 | **0.005** |
| K18578 | Cell wall degradation | 13 | 37 | 2 | 7 | 16 | 11 | 1 | 3 | 10 | 12 | 56 | 2 | 4 | 10 | 12 | <1 | <1 | 4 | **<0.001** |
| K18651 | Cell wall degradation | 12 | 41 | 2 | 6 | 19 | 7 | 1 | <1 | 12 | 10 | 60 | 2 | 4 | 16 | 4 | <1 | <1 | 4 | **<0.001** |
| K18786 | Cell wall degradation | 3 | 49 | 11 | 16 | | | 6 | <1 | 13 | 2 | 75 | 9 | 8 | | | 1 | | 5 | **<0.001** |
| K18576 | Cell wall degradation | | 14 | | 2 | 67 | 8 | <1 | | 8 | <1 | 24 | | **1** | 61 | 7 | <1 | | 7 | **<0.001** |
| K01202 | Primary metabolism | 3 | 16 | | 2 | 47 | 12 | <1 | 5 | 15 | 7 | 26 | | <1 | 47 | 10 | <1 | 2 | 8 | **<0.001** |
| K20455 | Primary metabolism | | 41 | 24 | 34 | | | | | | <1 | 48 | 21 | 31 | | | | | | **<0.001** |
| K02847 | Primary metabolism | 42 | 31 | 5 | <1 | | 19 | | | 2 | 36 | 39 | 9 | <1 | | 15 | | | 1 | **0.001** |
| K08961 | Primary metabolism | | | | | 3 | 30 | 1 | 54 | 12 | | | | | <1 | 67 | <1 | 30 | 3 | **<0.001** |
| K08325 | Primary metabolism | 3 | 28 | 11 | 16 | 3 | 17 | <1 | | 22 | 2 | 37 | 8 | 14 | 9 | 21 | | <1 | 9 | **<0.001** |
| K00211 | Primary metabolism | | | | | | 99 | | | 1 | | | | | | 100 | | | <1 | 0.201 |
| K03181 | Primary metabolism | | 89 | 11 | <1 | | | | | <1 | | 93 | 6 | <1 | | | | | <1 | 0.773 |
| K01085 | Primary metabolism | 55 | <1 | 9 | 11 | | | | | 24 | 84 | | 7 | 4 | | | | | 5 | **<0.001** |
| K01355 | Primary metabolism | 100 | | | <1 | | | | | | 99 | | <1 | <1 | | | | | <1 | 0.151 |
| K01819 | Primary metabolism | 47 | 18 | | 6 | 22 | | | | 7 | 59 | 24 | <1 | 8 | 7 | | | | 2 | **<0.001** |
| K16215 | Primary metabolism | | 2 | 25 | 10 | 56 | | | | 7 | | 3 | 11 | 3 | 81 | | | | 2 | **0.006** |
| K00998 | Primary metabolism | | 6 | 86 | 8 | | | | | | | <1 | 97 | 3 | | | | | <1 | NA |

| | | | | | | | | | | | | | | | | | | | | |
|---|---|---|---|---|---|---|---|---|---|---|---|---|---|---|---|---|---|---|---|---|
| K21280 | Primary metabolism | 80 | | | <1 | | | | | 19 | 99 | | | | | | | | <1 | **0.001** |
| K12455 | Primary metabolism | | 53 | | 14 | | 4 | | | 30 | | 87 | | 4 | | 2 | | | 7 | **<0.001** |
| K21239 | Primary metabolism | 100 | | | | | | | | | 100 | | | | | | | | | 0.151 |
| K00608 | Primary metabolism | | | | | | | | | 100 | | | | | | | | | 100 | **0.004** |
| K01352 | Primary metabolism | 11 | | | 3 | 29 | | | | 58 | 8 | | | 20 | 65 | | | | 7 | **0.004** |
| K02383 | Sensor regulator | | | | | | | | | 100 | | | | | | | | | 100 | 0.964 |
| K02399 | Sensor regulator | | 65 | 15 | 18 | | | | | 2 | | 81 | 14 | 5 | | | | | <1 | **0.009** |
| K02532 | Sensor regulator | 27 | 45 | | 12 | 3 | | 12 | | 1 | 23 | 67 | <1 | 6 | <1 | | 3 | | 1 | **<0.001** |
| K03202 | Sensor regulator | 96 | | | 4 | | | | | <1 | 99 | | | <1 | | | | | <1 | 0.150 |
| K03397 | Sensor regulator | | 14 | | | | | | | 86 | | 77 | | 3 | | | | | 20 | **0.010** |
| K03477 | Sensor regulator | 58 | 10 | 4 | 3 | 5 | | | 2 | 19 | 58 | 29 | 1 | <1 | 4 | <1 | | <1 | 8 | **<0.001** |
| K04338 | Membrane transport | 43 | 39 | 3 | 10 | | 2 | | | 3 | 46 | 25 | 3 | 4 | 8 | 14 | | | <1 | **0.018** |
| K04643 | Membrane transport | 33 | | | 21 | 6 | 18 | | | 22 | 14 | | | 1 | 6 | 77 | | | 2 | **0.001** |
| K05372 | Membrane transport | 4 | | 10 | 12 | | 57 | 1 | | 16 | 17 | | 2 | 4 | | 68 | <1 | | 8 | **<0.001** |
| K05660 | Membrane transport | | | | | | | | | 100 | | | | | | | | | 100 | 0.379 |
| K05874 | Membrane transport | 10 | 68 | 4 | 7 | <1 | <1 | <1 | <1 | 11 | 14 | 74 | 5 | 4 | <1 | <1 | <1 | | 3 | **<0.001** |
| K05876 | Membrane transport | | 93 | <1 | 6 | | | | | 1 | | 96 | 1 | 3 | | | | | | **0.004** |
| K06080 | Membrane transport | | | | | | | | | 100 | | | | | | 89 | | | 11 | **<0.001** |
| K07345 | Membrane transport | | 61 | 30 | 9 | | | | | | | 48 | 47 | 4 | | | | | <1 | 1.000 |
| K07786 | Membrane transport | | | | | 73 | 13 | | | 14 | | | | | 37 | 60 | | | 3 | **<0.001** |
| K07991 | Membrane transport | | | | | | | | | 100 | | | | | | | | | 100 | **0.001** |
| K08156 | Membrane transport | | 38 | 2 | <1 | | 57 | | | 2 | | 61 | 1 | <1 | <1 | 36 | | | 2 | **0.003** |
| K08566 | Membrane transport | 77 | | | 3 | | | | | 20 | 98 | | | 2 | | | | | | **0.009** |
| K10094 | Membrane transport | 6 | 76 | 2 | 15 | | | | | 1 | 4 | 88 | 3 | 5 | | | | | <1 | **0.006** |

| | | | | | | | | | | | | | | | | | | |
|---|---|---|---|---|---|---|---|---|---|---|---|---|---|---|---|---|---|---|
| K10235 | Membrane transport | 67 | 20 | | 8 | 1 | | | 4 | 72 | 22 | <1 | 4 | <1 | | | 2 | **<0.001** |
| K10857 | Membrane transport | 85 | 1 | <1 | 2 | 5 | | | 6 | 93 | <1 | <1 | 1 | 4 | | | 2 | **<0.001** |
| K10926 | Biofilm formation | | 84 | 8 | 3 | | | | 5 | | 94 | 4 | <1 | | | | 1 | **<0.001** |
| K10927 | Biofilm formation | | 87 | 7 | 2 | | | | 4 | | 95 | 4 | <1 | | | | 1 | **<0.001** |
| K10975 | Biofilm formation | | | | | 56 | | 14 | 30 | | | | | 100 | | | <1 | **0.005** |
| K11017 | Biofilm formation | 18 | 61 | 14 | 8 | | | | | 28 | 60 | 10 | 2 | | | | <1 | 0.536 |
| K11734 | Biofilm formation | | 39 | 11 | 4 | 1 | 1 | 12 | 32 | <1 | 64 | 10 | 2 | <1 | 7 | 5 | 13 | **<0.001** |
| K11743 | Biofilm formation | | | 21 | 13 | | | | 66 | 22 | | 59 | 6 | <1 | | | 12 | **0.001** |
| K11889 | Biofilm formation | 88 | | | <1 | | | | 12 | 100 | | | <1 | | | | <1 | **0.002** |
| K11909 | Biofilm formation | 66 | | 7 | 24 | | | | 3 | 88 | | 3 | 7 | | | | 3 | **0.005** |
| K11934 | Quorum sensing | | | | 4 | | 95 | | 1 | - | | | <1 | | 99 | | <1 | **<0.001** |
| K11964 | Quorum sensing | 70 | | | | | | | 30 | 100 | | | | | | | | 0.054 |
| K12059 | Quorum sensing | 54 | 40 | 3 | 3 | | | | <1 | 66 | 32 | <1 | <1 | | | | 1 | **<0.001** |
| K12069 | Quorum sensing | 69 | 21 | | 5 | | | | 5 | 75 | 22 | | 2 | | | | 1 | **<0.001** |
| K12083 | Quorum sensing | | 26 | | 51 | | | | 23 | | 84 | | 10 | | | | 6 | **0.014** |
| K12228 | Quorum sensing | | 17 | | 33 | | | | 50 | | 86 | | 14 | | | | | **0.006** |
| K12280 | Transcriptional regulation | | 87 | 12 | <1 | | | | <1 | | 95 | 5 | <1 | | | | <1 | 0.965 |
| K12281 | Transcriptional regulation | | 93 | 2 | 5 | | | | | | 95 | 5 | <1 | | | | <1 | NA |
| K12284 | Transcriptional regulation | | 94 | 5 | <1 | | | | | | 93 | 7 | <1 | | | | <1 | NA |
| K12285 | Transcriptional regulation | | 90 | 7 | 2 | | | | | | 96 | 4 | <1 | | | | <1 | NA |
| K12286 | Transcriptional regulation | | 93 | 5 | 2 | | | | | | 95 | 5 | | | | | | NA |
| K12688 | Secretion systems | 3 | 13 | 79 | 5 | | | | <1 | 2 | 4 | 93 | <1 | | | | 1 | 0.145 |
| K12820 | Secretion systems | | | | | | | | 100 | | | | | | | | 100 | **<0.001** |
| K13448 | Secretion systems | | | | | 69 | | | 31 | | | | | 99 | | | <1 | **0.033** |

| | | | | | | | | | | | | | | | | | | | | |
|---|---|---|---|---|---|---|---|---|---|---|---|---|---|---|---|---|---|---|---|---|
| K13454 | Secretion systems | | | | | | | | | | | 44 | | | | | | | <u>56</u> | NA |
| K13585 | Secretion systems | <u>90</u> | | | | | | | | 10 | <u>92</u> | | | | | | | | 8 | 0.050 |
| K13586 | Secretion systems | <u>94</u> | <1 | | 4 | | | | | <1 | <u>97</u> | <1 | | 2 | | | | | <1 | 0.134 |
| K13831 | Secretion systems | | | | | | | | | <u>100</u> | | | | | | | | | <u>100</u> | **<0.001** |
| K13964 | Secretion systems | | | | | <u>98</u> | | | | 2 | | | | | <u>99</u> | | | | 1 | 0.466 |
| K14781 | Secretion systems | | | | | | | | | <u>100</u> | | | | | | | | | <u>100</u> | **<0.001** |
| K16088 | Secretion systems | <u>42</u> | 27 | 23 | 6 | | | | <1 | 1 | <u>42</u> | 30 | 25 | 2 | | | | <1 | 1 | 0.057 |
| K16210 | Secretion systems | <u>55</u> | 8 | 3 | 13 | | | 7 | | 14 | <u>62</u> | 23 | 2 | 8 | | | 2 | | 3 | **<0.001** |
| K16347 | Secretion systems | <u>41</u> | 17 | | 31 | | | | | 11 | <u>56</u> | 42 | | 3 | | | | | | **0.003** |
| K16348 | Secretion systems | <u>85</u> | 6 | | 9 | | | | | | <u>88</u> | 11 | | <1 | | | | | <1 | 0.151 |
| K16552 | Secretion systems | <u>100</u> | | | <1 | | | | | <1 | <u>100</u> | | | | | | | | <1 | 0.964 |
| K16553 | Antibiotic resistance | <u>59</u> | | | <1 | 4 | | | | 36 | <u>89</u> | | | <1 | 3 | | | | 8 | **<0.001** |
| K16696 | Antibiotic resistance | | | 11 | 11 | | | | | 78 | | | 91 | 9 | | | | | <1 | **<0.001** |
| K17060 | Antibiotic resistance | 16 | <u>74</u> | 2 | 4 | | | | | 3 | 14 | <u>81</u> | 2 | 2 | | | | | 1 | **<0.001** |
| K18373 | Antibiotic resistance | | <u>99</u> | | <1 | | | | | <1 | | <u>99</u> | | <1 | | | | | <1 | 0.754 |
| K18380 | Antibiotic resistance | | <u>99</u> | | <1 | | | | | <1 | | <u>100</u> | | <1 | | | | | <1 | 0.965 |
| K18642 | Motility | <u>100</u> | | | | | | | | | <u>100</u> | | | | | | | | | NA |
| K18793 | Motility | | | | | | <u>92</u> | | | 8 | | <1 | | <1 | | <u>99</u> | | | <1 | **<0.001** |
| K19033 | Secondary messaging | | | 23 | | | <u>74</u> | | | 3 | | | 40 | <1 | | <u>59</u> | | | <1 | **0.002** |
| K19060 | Secondary messaging | <u>65</u> | | | <1 | 18 | | | | 16 | <u>76</u> | | | 2 | 17 | | | | 5 | **<0.001** |
| K19101 | Defence | 8 | 32 | 1 | <u>45</u> | | | | | 14 | 9 | <u>59</u> | 1 | 25 | | | | | 5 | **<0.001** |
| K19213 | Defence | 45 | | | <u>54</u> | | | | | 1 | <u>74</u> | 17 | | 9 | | | | | <1 | 0.151 |
| K19216 | Defence | | | | | | 94 | | | 6 | | | | | | <u>99</u> | | | 1 | **<0.001** |
| K19611 | Defence | | | <u>63</u> | 3 | | | | | 34 | | | <u>98</u> | 2 | | | | | <1 | **0.002** |
| K19731 | Calcium sensors | <u>67</u> | 24 | 3 | 2 | | | | | 4 | <u>66</u> | 32 | <1 | 1 | | | | | 1 | **0.003** |

| | | | | | | | | | | | | | | | | | | | | |
|---|---|---|---|---|---|---|---|---|---|---|---|---|---|---|---|---|---|---|---|---|
| K20267 | Cell cycle | 99 | | | 1 | | | | | | 100 | | | | | | | | | **<0.001** |
| K20268 | Cell cycle | 17 | 4 | 13 | 7 | 25 | 5 | | | 29 | 13 | 12 | 12 | 4 | 17 | 16 | | | 26 | **0.050** |
| K20272 | Cell cycle | 90 | | | | | | | | 10 | 100 | | | | | | | | | 0.117 |
| K20275 | Cell cycle | | | 10 | 11 | | 51 | | | 28 | 3 | | 8 | 6 | | 70 | | | 14 | **<0.001** |
| K20326 | Cell cycle | 42 | | | 2 | | 55 | | | <1 | 51 | | | <1 | | 48 | | | 1 | **0.010** |
| K20555 | Primary metabolism | 60 | | | | | | | | 40 | 90 | - | | | | | | | 10 | 0.076 |
| K20961 | Primary metabolism | | 72 | 5 | 14 | <1 | | | | 9 | - | 86 | 3 | 7 | | | | | 3 | **<0.001** |
| K20966 | Primary metabolism | 56 | | | 31 | | | | | 13 | 78 | | 2 | 14 | | | | | 6 | **<0.001** |
| K21346 | Membrane transport | | | | | | | | | 100 | | | | | | | | | 100 | **<0.001** |
| K21825 | Motility | 35 | 48 | 5 | 2 | 2 | | | 7 | <1 | 30 | 60 | 5 | <1 | 3 | | <1 | 2 | 1 | **<0.001** |
| K22335 | Motility | 49 | 12 | | 37 | | | | | 1 | 61 | 19 | | 19 | | | | | 1 | 0.773 |
